# A Synthetic Virology Platform for Rapid Prototyping of Influenza A Virus mutants

**DOI:** 10.64898/2026.10.24.761556

**Authors:** Janarthan Rama Murti, Karin Schön, Davide Angeletti

## Abstract

Influenza A virus (IAV) remains a significant public health threat, with the need for effective vaccines and antiviral therapies remaining a high priority. *In vitro* testing is paramount in studying novel IAV viruses and their immunogenicity. To address this need, we developed a synthetic virology approach to rapidly prototype and test IAV mutants with altered antigenic sites. By strategically inserting restriction sites within an ambisense plasmid, we generated a tool where we can seamlessly replace either 1) the full hemagglutinin (HA), 2) the HA globular head domain to generate head/stalk chimeras or 3) the five canonical antigenic sites. The cloning inserts can be computationally designed and artificially synthesized, enabling rescue of synthetic Influenza A viruses. Using this platform we rescued IAV H1N1, H3N2 & H5N1, and went on to prototype their respective antigenic site mutants & chimeric IAVs using viral genome sequence alone. This platform could be useful for generating vaccine seed strains or to study anti-viral immunity.

Figure 0.
Abstract Figure

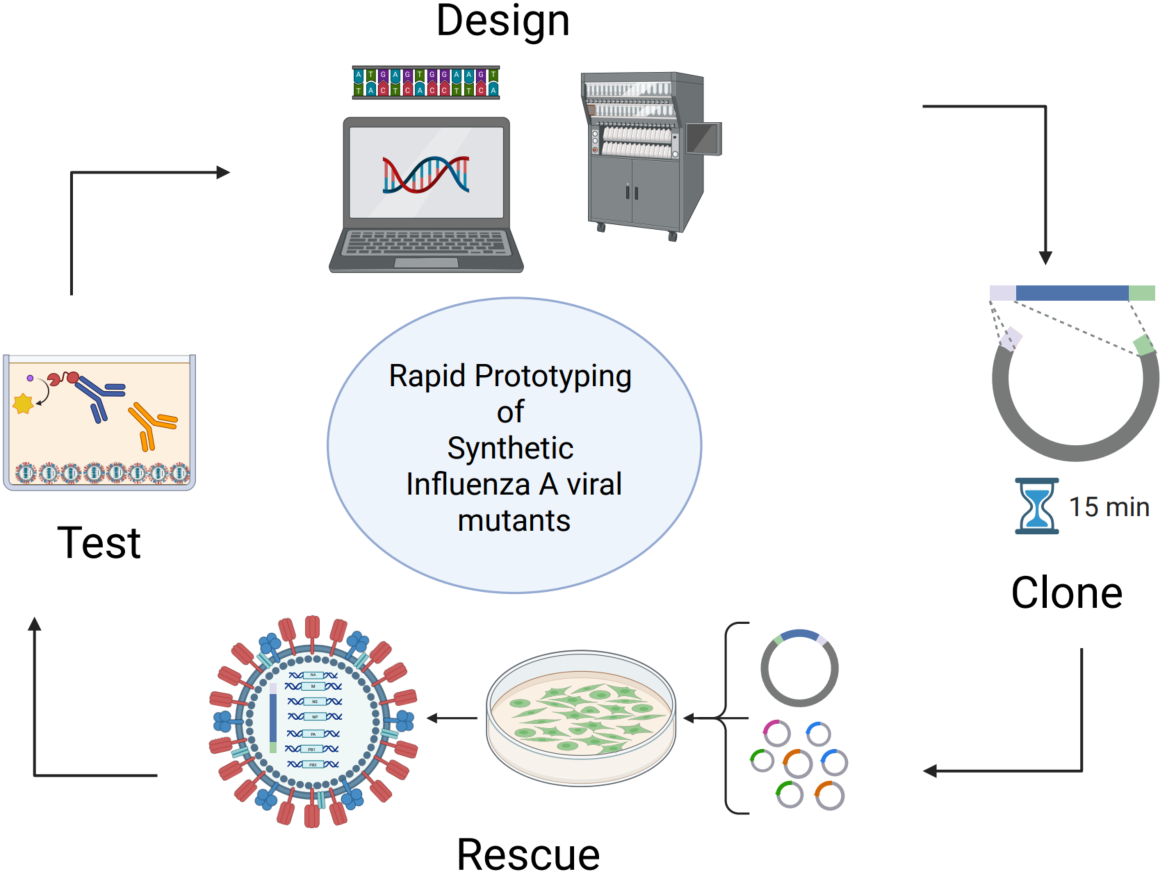

## Introduction

Influenza A virus (IAV) is a significant public health concern, causing seasonal epidemics and occasional pandemics worldwide (Al Hajjar & McIntosh, 2010; Hadfield et al., 2018; Shu & McCauley, 2017a; Tübingen, 2009). The development of effective vaccines and antiviral therapies is essential for controlling the spread of influenza viruses. However, the emergence of viral variants that escape host immune responses poses a significant challenge to the development of such interventions.

Reverse genetic has revolutionized IAV research by allowing for the manipulation of viral genomes to create recombinant RNA viruses via specific genetic modifications done on viral cDNA (Enami & Palese, 1991; Hoffmann et al., 2000; Luytjes et al., 1989; Neumann et al., 1999). Synthetic virology, on the other hand, has emerged as a powerful approach to design and engineer synthetic viral genomes from chemically synthesized DNA (Allen & Iii, 2021; Benjamin, 2023). Assembly of viral genetic material using synthetic genes or even entire virions in the lab has been made possible by artificial DNA synthesis technology (Cello et al., 2002; Dormitzer et al., 2013; Noyce et al., 2018). Here, we propose a platform that puts together synthetic virology and reverse genetics approaches to design, rescue, and characterize IAV mutants.

IAV antigenic site mutants are indispensable tools to explore mutational spaces and trajectories of HA antigenic sites and to characterize antiviral immune responses. However, there is a barrier in rescuing such mutants and is often the most technically challenging step in conducting mutational antigenic profiling and immunodominance studies (Bloom, 2014; Broecker et al., 2018; Burton & Eyre, 2021; Doud et al., 2017; Lee et al., 2018; S. T. H. Liu et al., 2018; Martínez et al., 2024). Current technologies, such as PCR-amplification/mutagenesis of viral genome segments, multi-segmental ligation and mutagenesis by in vitro CRISPR-Cas9 have technical and scalability challenges and require virus isolate-derived genetic templates. These methods are often poorly scalable, especially when creating libraries of multi-antigenic site IAV mutants via saturation mutagenesis and for the prototyping of mosaic IAV mutant viruses (Markushin & Rtishchev, 2017; Ye et al., 2015).

To overcome these limitations, we describe a methodological framework for a plasmid-based synthetic virology platform that enables the rapid design and rescue of synthetic IAV mutants (Figure 1). As proof of concept, we used this platform for the generation of i) IAV Hemagglutinin (HA) subtypes, ii) HA antigenic site mutants and iii) HA Head/stalk chimera solely using viral genomic sequences available in public databases as our input material. Our approach will facilitate the generation of novel viruses that could be used for immunological or vaccine studies

**Figure 1.**
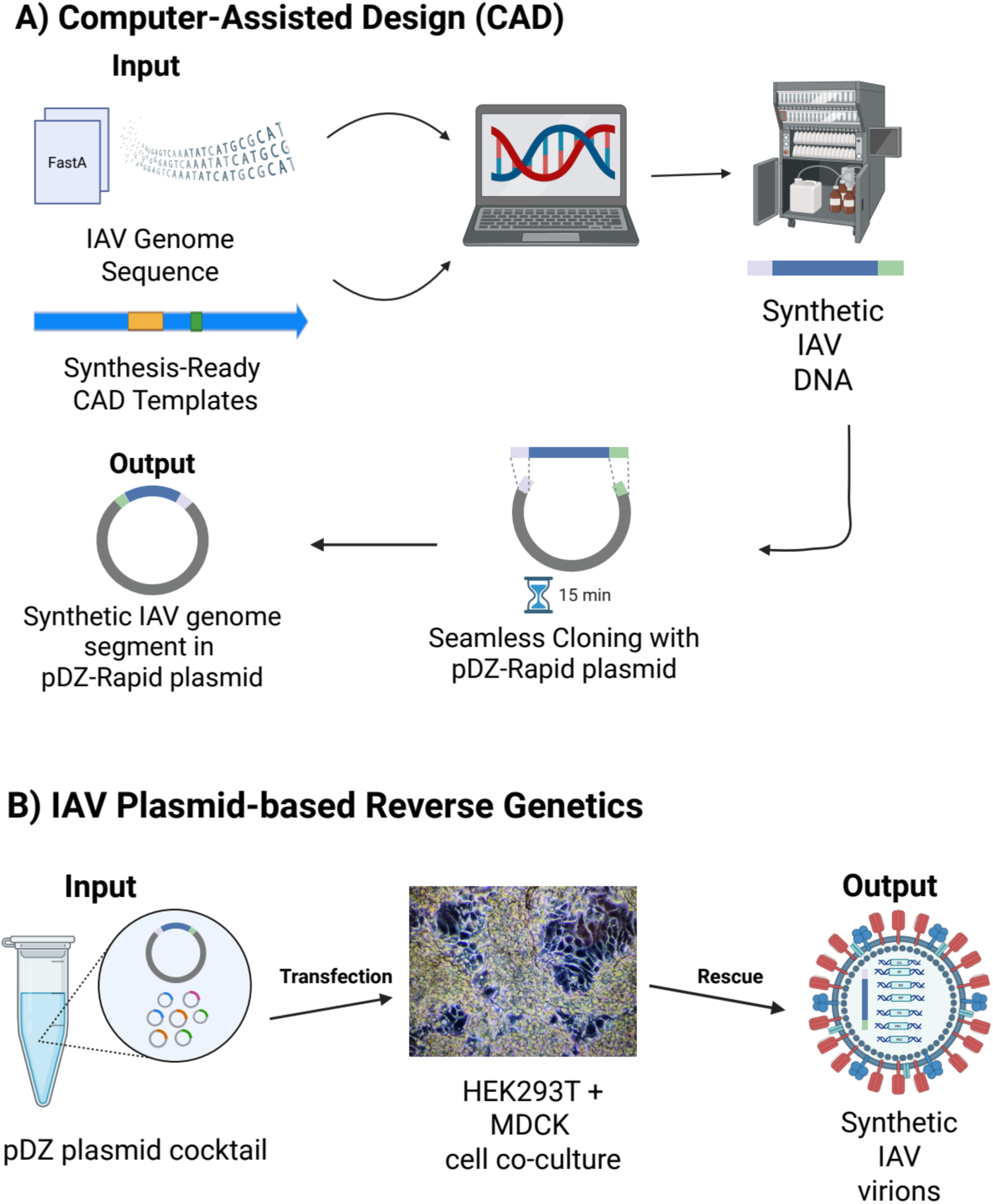
Introduction figure, overview of methodological framework to prototype synthetic Influenza A virus (IAV). A) Synthetic IAVs are designed with Computer Assisted Design (CAD) templates and using published IAV genome sequence as input. The designed construct is synthesized and seamlessly cloned into the pDZ-Rapid plasmid, without requiring a physical template. B) A pDZ plasmid cocktail is made with the cloned pDZ plasmid and pDZ plasmids containing the appropriate backbone IAV genome segments. This pDZ plasmid cocktail is transfected into a co-culture of HEK293T & MDCK cells to produce virions of the designed synthetic IAV

## Results

### Modification of an ambisense plasmid for rapid H1 ORF exchange

To overcome current limitations and cumbersome cloning strategies inherent to the pDZ plasmid and to be able to insert and modify the HA open reading frame (ORF) and the UTRs exclusively, we decided to insert four novel features to existing pDZ plasmid, while maintaining its ambisense properties (Hoffmann et al., 2000). The first feature is ability to prototype IAV of any HA subtype (HAx ORF cloning). The second is to generate antigenic site mutants, via antigenic site cassette mutagenesis (ASC), the third is to produce head/stalk chimera (GHD cloning and the fourth is to enable site-directed saturation mutagenesis.

To achieve our goals and to identify strategically placed conserved restriction sites for the removal of the Antigenic Site Cassette (ACS) and Globular Head Domain (GHD) we aligned 5877 H1N1 HA ORF sequences spanning from 1991 to 2025 and identified HpaI, BstBI and MscI sites as highly conserved and appropriately localized (Figure 2A-B).

**Figure 2.**
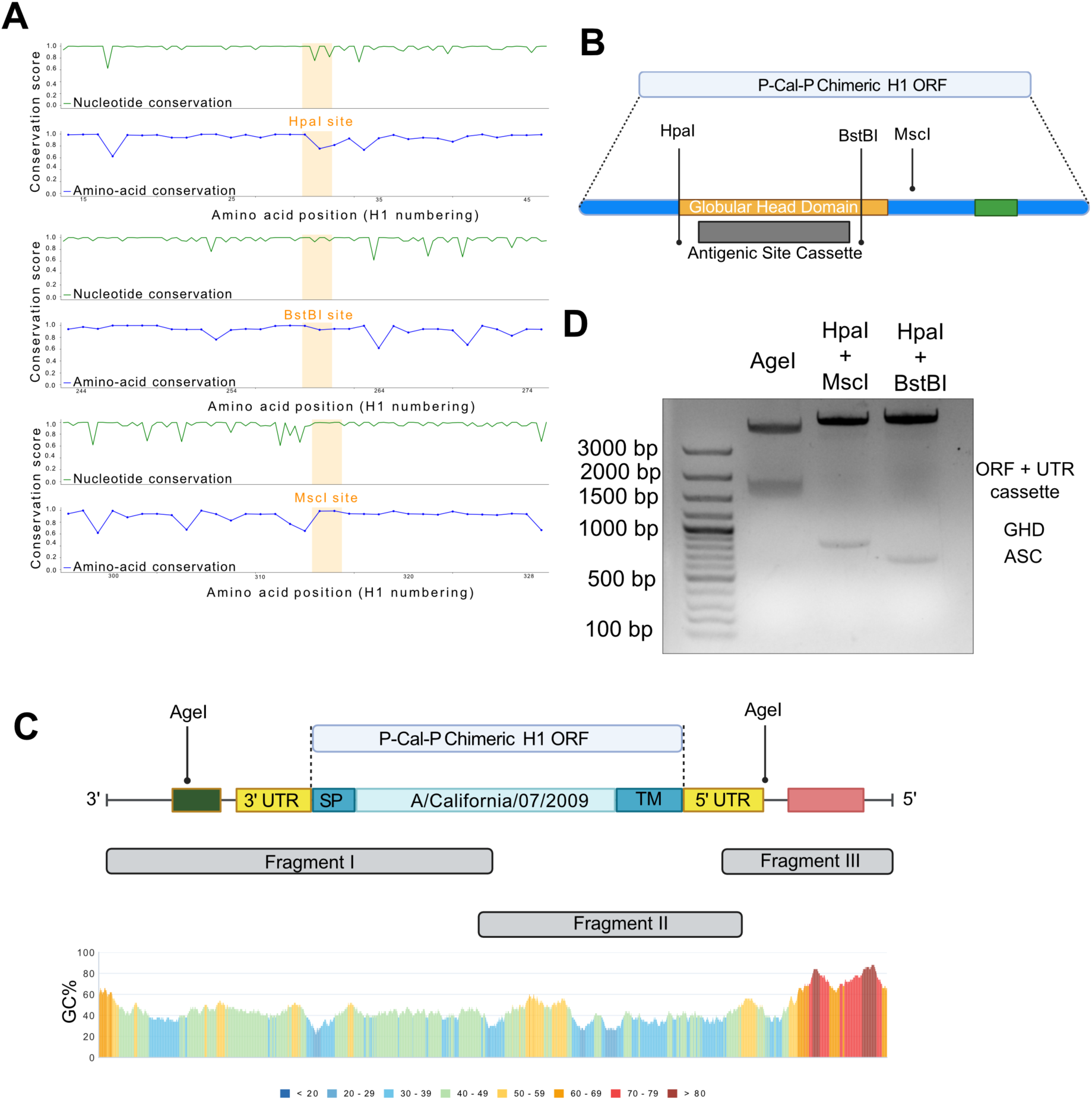
Creation of pDZ-Rapid plasmid. A) Restriction site conservation analysis of the HpaI, BstBI and MscI restriction sites in H1N1 IAV. The nucleotide and amino acid conservation tracings are shown in green and blue respectively. The H1 numbering system has been used for the amino acid sequence. B) Strategic location of HpaI, BstBI and MscI restriction sites in the P-Cal-P pdmH1 ORF. The P-Cal-P pdmH1 ORF (blue) contains globular head domain (GHD, orange), Antigenic Site Cassette (ASC, grey) and cleavage site (green). C) Complete assembly of Fragment I, II and III with aligned GC content tracing. The labelled restrictions sites were inserted to flank the P-Cal-pdmH1 ORF and its UTRs. fragments I, II and III were designed to have 20 bp overlap sequences, to enable directional seamless cloning into the linearized pDZ backbone. The fragments contain the following genetic parts: I) Fragment I contains the mouse RNA polymerase I terminator (dark green), viral 3’ UTR (yellow) and part of the P-Cal-P chimeric ORF. The P-Cal-P H1 chimeric ORF in fragment I contains the PR8 signal peptide (SP), the 5’ half of the A/California/07/2009 (pdmH1N1) ORF. This fragment is 894 bp, contains all five H1 canonical antigenic sites, and has a GC content of 43%. II) Fragment II contains the 3’ half of the A/California/07/2009 (pdmH1N1) H1 ORF, the PR8 transmembrane region and cytoplasmic tail (TM), and the viral 5’ UTR (yellow). Fragment II is 971 bp and has 40% GC content. III) Fragment III contains the human RNA polymerase I promoter (hRNAP promoter, red), is 314 bp and has 71% GC content. D) Restriction analysis of the final pDZ-Rapid plasmid assembly. Agarose gel image containing bands (from left to right) of the pDZ-Rapid plasmid digested for HAx ORF cloning, HA head-stalk chimera cloning and HA ASC mutagenesis respectively using the three restriction enzyme combinations indicated in the figure.

These were chosen in particular as they were strategically placed and importantly their insertion did not require making non-synonymous mutations on the pdmH1 ORFs. For a deeper insight, we re-analyzed the conservation of the identified restriction sites using a small curated set of IAV H1 sequences ranging from 1918 to 2022 (Figure S1), confirming their amino acid conservation, despite some variation at the nucleotide level. Indeed, the human seasonal pdmH1N1 IAV strains lost their HpaI site by one nucleotide change, since 2019: still, this could be reintroduced via a synonymous mutation (Figure S1A).

We derived our plasmid backbone from a pDZ plasmid linearized with EcoRI and XhoI restriction enzymes (Figure S2A) and inserted a designed chimeric HA ORF. This ORF (P-Cal-P H1) derives its HA signal peptide and transmembrane region sequences (P) from the lab-adapted PR8 H1N1 strain (A. R. Taylor et al., 1943), while the bulk of the H1 ORF originates from Influenza A/California/07/2009 (pdmH1N1) (Kawahara et al., 2020). Finally, we inserted an AgeI site proximal to the 3’ UTR to compliment the existing AgeI near the 5’ UTR. In this way we have two AgeI restrictions sites flanking the HA ORF at its 5’ and 3’ UTRs (Figure 2C).

The construct of 2103 bp containing the P-Cal-P H1 ORF and its flanking genetic parts was designed as three overlapping oligonucleotide fragments, i.e, Fragment I, II, and III (Figure 2C). This construct was divided into the three fragments to meet the design constraints, including size and GC content. Although fragments I and II were successfully synthesized using solid phase phosphoramidite reaction cycle, fragment III could not be synthesized due to its high GC content (Figure 2C). We instead PCR amplified fragment III from a template pDZ plasmid to have 20 bp overlapping regions with fragment II and the pDZ plasmid backbone (Figure S2B). We then cloned these three overlapping insert fragments using the NEBuilder HiFi DNA Assembly technique into the linearized plasmid backbone (Figure S2A), to form the final assembly, named pDZ-Rapid plasmid.

In summary, the generated pDZ-Rapid plasmid included the following restriction sites: 5’ AgeI, HpaI, BstBI, MscI, and 3’ AgeI (Figure 2C). By linearizing the pDZ-Rapid plasmid using different combinations of enzymes, we could manipulate the Influenza A H1 structure to remove and exhange the HA ORF and its flanking UTRs (AgeI digestion; Figure 2B-D), the HA globular head domain (GHD) (HpaI, MscI digestion; Figure 2B-D), or the HA antigenic site cassette (ASC) (HpaI, BstBI digestion; Figure 2B-D). Restriction analyses with the respective restriction enzyme combinations confirmed their compatibility for efficient double digest and the correct final assembly (Figure 2D). In addition, design of inserts is facilitated by the usage of Computer-Aided Design (CAD) templates (see Supplementary Files, SF1 - SF3).

Overall, the pDZ-Rapid plasmid is configured for rapid prototyping of synthetic IAV mutants and chimeric viruses.

### Rescue of hybrid Influenza A P-Cal-P pdmH1: PR8 with assembled pDZ-Rapid Plasmid

To validate the functionality of the pDZ-Rapid plasmid, we rescued the P-Cal-P pdmH1 IAV as a hybrid using the laboratory-adapted PR8 backbone. To this end, we co-transfected the HA from P-Cal-P pdmH1 pDZ-Rapid plasmid with 7 other PR8 pDZ plasmids i.e., NA, NP, M, NS, PA, PB1, PB2 segments (Hoffmann et al., 2002; Webster, 1970).

We demonstrated the rescue of hybrid P-Cal-P pdmH1: PR8 IAV via hemagglutination assay (Figure 3A), confirming that the modifications we made did not affect plasmid functionality. The hybrid P-Cal-P pdmH1: PR8 IAV grew well and demonstrated higher fitness than the wild type A/California/07/2009 (pdmH1N1) (Fig 3B), possibly owing to the flanking PR8 regions inserted. Finally, we used the S-OIV 5H7 mAb, recognizing only pdmH1N1 but not classical swine H1N1 strains (Shao et al., 2011) and demonstrated antigenic recognition of P-Cal-P pdmH1: PR8 virus but not wild type PR8 virus (Figure 3C). In conclusion, we confirmed that the pDZ-Rapid plasmid is functional and the rescued virus is functional.

**Figure 3.**
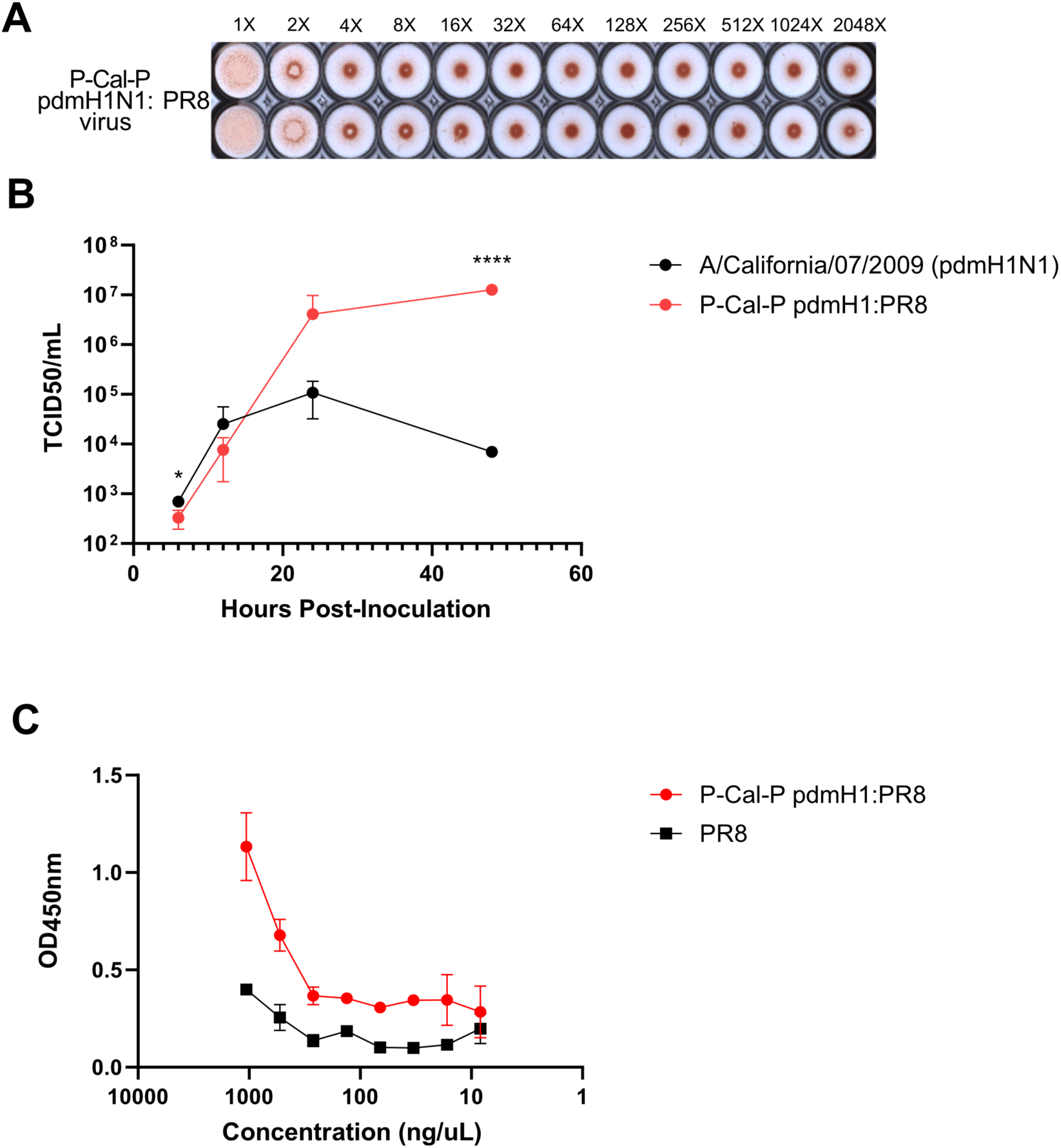
Rescue and characterization of P-Cal-P pdmH1: PR8 IAV. A) Hemagglutination assay of the P-Cal-P pdmH1: PR8 virus rescued via transfection of the newly constructed pDZ-Rapid plasmid using PR8 backbone. B) Growth curve of the rescued P-Cal-P pdmH1: PR8 IAV compared to wild type A/California/07/2009 (pdmH1N1). Two-way ANOVA multiple comparison * p-value < 0.05 at 6 hours post-inoculation and **** p-value < 0.0001 at 48 hours post-inoculation. C) Antigenic characterization of rescued P-Cal-P pdmH1: PR8 IAV against S-OIV 5H7 monoclonal antibody (mAb) compared to wild type PR8 H1N1 IAV as control. Both IAVs were normalized with hemagglutination assay, and 4 hemagglutinating units of both viruses were used for ELISA testing.

### Creation of PR8-like H1N1 variant of the pDZ-Rapid via Rapid HA ORF exchange

Next, we set out to create a PR8 H1 variant of the pDZ-Rapid plasmid while maintaining the same features described previously, as a proof of concept but also as the PR8 H1N1 is the most widely used laboratory-adapted strain for basic IAV research *(A. R. Taylor et al., 1943; R. M. Taylor & Dreguss, 1941)*. While the PR8 H1 sequence maintains the HpaI restriction site, this virus did not have the BstBI restriction site. We therefore inserted the BstBI restriction site at position 255 (H1 numbering), also introducing an A255F mutation (Figure 4A). Likewise, the MscI site was not conserved in PR8 (Figure S1c) but the GHD could be removed via HpaI and DraIII double digestion (Figure 4B)

**Figure 4.**
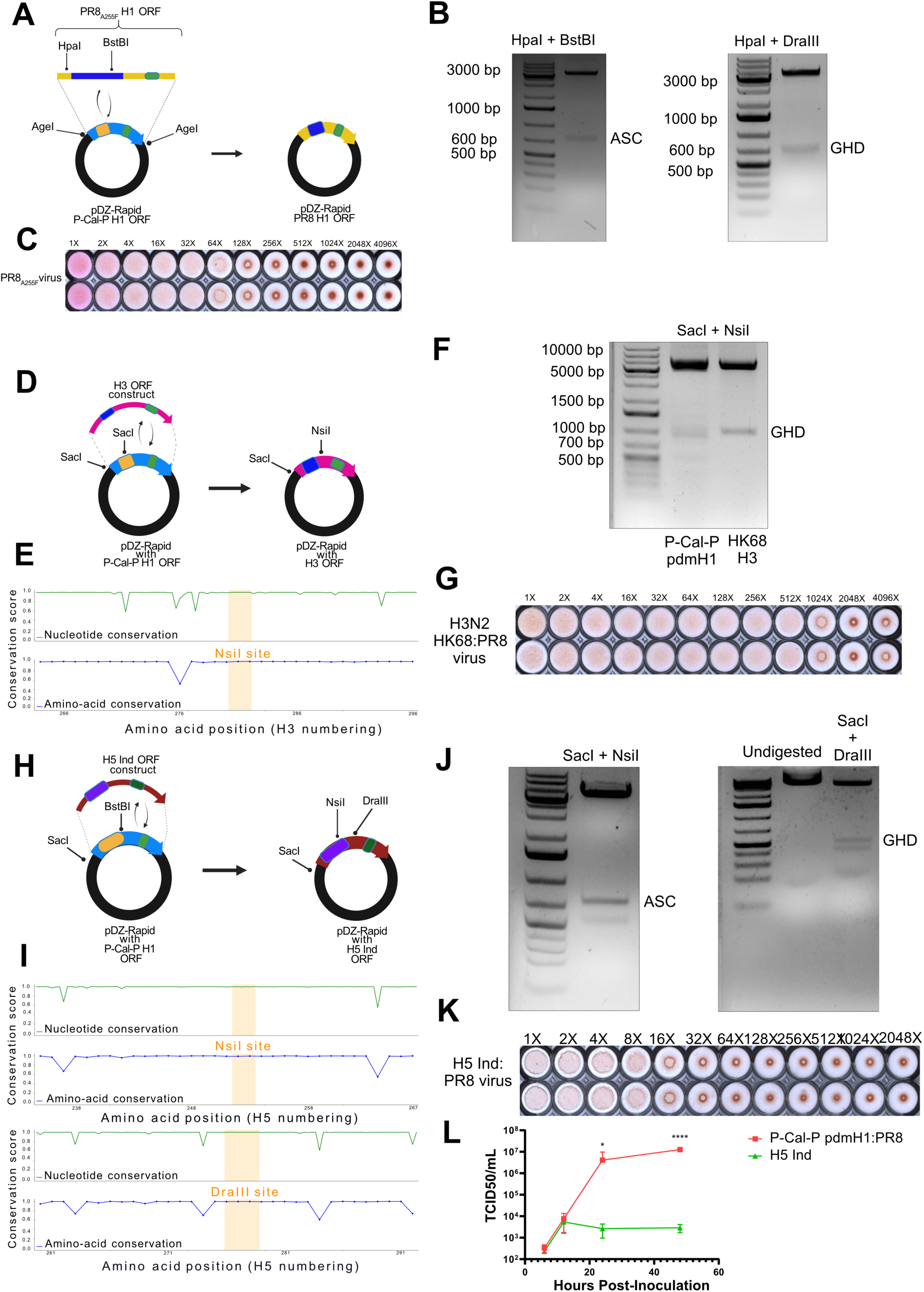
Creation of pDZ-Rapid plasmid variants for prototyping different IAV HA subtypes via digestion by AgeI restriction enzyme. A) Creation of PR8 variant of pDZ-Rapid plasmid. The PR8 A255F H1 ORF (yellow, with PR8 Antigenic Site Cassette (ASC) in dark blue and HA cleavage site in green) was synthesized and seamlessly cloned into the pDZ-Rapid plasmid linearized by AgeI. B) Gel image with HpaI + BstBI double digestion of pDZ-Rapid PR8 variant, resulting in a restriction fragment with all five canonical antigenic sites of PR8 H1 (left). Gel image of HpaI & DraIII digestion of pDZ-Rapid PR8 variant, producing a restriction fragment with the PR8 H1 globular head domain (GHD) (right). C) Hemagglutination assay of the rescued PR8 IAV. D) Creation of H3 variant of pDZ Rapid plasmid. A synthesized single fragment H3 ORF of HK68 IAV (pink, with H3 GHD in dark blue and HA cleavage site in green) was seamlessly cloned into the pDZ Rapid plasmid containing a SacI restriction site in the plasmid backbone. The cloned plasmid ended up with a unique SacI site which is useful to remove the H3 GHD via double digest with SacI + NsiI restriction enzymes. E) Restriction site conservation analysis of NsiI site in H3N2 IAV. F) Gel image of restriction digest of pDZ Rapid P-Cal-P and pDZ Rapid H3 HK68 with SacI + NsiI restriction enzymes. G) Hemagglutination assay of HK68 H3N2 IAV rescued with PR8 backbone. H) Creation of H5 variant of pDZ-Rapid plasmid. The H5 ORF of A/Indonesia/05/2005/H5N1 (H5 Ind) was computationally attenuated via conversion of its polybasic cleavage site into a bi-basic cleavage site, synthesized and then cloned into pDZ Rapid plasmid. The H5 ORF is shown in red with its GHD in violet and the bi-basic cleavage site in dark green. I) NsiI and DraIII restriction site conservation analysis. J) Gel images of restriction digest of pDZ Rapid H5 with SacI + DraIII (right) and SacI + NsiI (left). K) Hemagglutination assay of the rescued H5 Ind: PR8 IAV. L) Growth curve characterization of the H5 Ind: PR8 IAV, in comparison with P-Cal-P pdmH1:PR8 virus (also shown in Figure 3B). Two-way ANOVA multiple comparison * p-value < 0.05 at 24 hours post-inoculation and **** p-value < 0.0001 at 48 hours post-inoculation.

Reverse genetics with this cloned PR8 H1 pDZ-Rapid plasmid enabled the rescue of a PR8 virus carrying the A255F mutation, which was able to induce hemagglutination (Figure 4C). While we attempted to simultaneously insert the BstBI and MscI sites via A255F and V314A mutations respectively, this non-synonymous mutation combination rendered the virus unviable.

### Creation of H3 variants of the pDZ-Rapid plasmid via Rapid HA ORF exchange

Next, we set out to create pDZ-Rapid plasmids for group 2 IAVs and generated a plasmid containing H3 of the HK68 H3N2 strain, which is a laboratory-adapted group 2 IAV strain.

To facilitate cassette mutagenesis of the H3 ORF, we modified fragment I by inserting a SacI site in the 5’ end of fragment I (Figure 4D). Cloning of H3 ORF into the pDZ-Rapid plasmid with the modified fragemnt I turned above mentioned SacI site into a unique restriction site. In the vicinity of the H3 GHD’s 3’ end, we identified an NsiI site as remarkably well conserved (Figure 4E). A deeper analysis corroborated the conservation of this NsiI restriction site for over more than 50 years of human seasonal H3N2 IAV evolution (Figure S3A). Hence restriction double digest of the H3 pDZ-Rapid plasmid with SacI and NsiI gave a clear single band containg the H3 GHD, as the H3 ORF has the unique NsiI site and no SacI site (Figure 4F, right lane). On the other hand, the same digestion of the P-Cal-P pdmH1 pDZ-Rapid plasmid gave mutiple bands (Figure 4F, left lane) as this plasmid contains another SacI restriction site within the P-Cal-P pdmH1 ORF (shown earlier in Figure 4D).

We successfully rescued the hybrid HK68 H3N2 virus by co-transfecting the HK68 H3 pDZ-Rapid plasmid with pDZ plasmids carrying N2 of HK68 strain and the PR8 backbone. Again, the virus induced hemagglutination (Figure 4G), grew well in cell culture (Figure S3B) and was recognized by AX-LAH3, a pan-H3 mAb (Figure S3C) and by anti-HK68 H3N2 mice polyclonal sera (Figure S3D).

### Creation of an attenuated H5 variant of the pDZ-Rapid plasmid via Rapid HA ORF exchange

As a last ORF-switch example, we explored the utility of pDZ-Rapid plasmid rapid prototyping of the H5 IAV subtype, the same subtype that is causing ongoing public health concerns (World Health Organization, 2024). Via restriction site conservation analysis of H5 ORF sequences, we identify two strategically placed restriction sites downstream the H5 ASC and GHD i.e., NsiI and DraIII, to enable removal of the H5 ASC and H5 GHD respectively (Figure 4H-I). Most H5 IAV strains including the recent North American clade 2.3.4.4b (Kandeil et al., 2023; Tian et al., 2023) carry well conserved amino acid sequences at the NsiI & DraIII sites (Figure 4I & S4).

An attenuated H5 ORF of Influenza A/Indonesia/05/2005 (H5N1) (Kandun et al., 2006) was designed by modifying it’s polybasic cleavage site and cloned into an AgeI-digested pDZ-Rapid plasmid (Figure 4J). Together with the SacI site inserted into fragment I earlier (Figure 4D), we configured the pDZ-Rapid plasmid for H5 ASC and GHD mutagenesis via the combinations of restriction enzymes mentioned above (Figure 4H-J). As a SacI site is present in the H5 ASC, in both the SacI + NsiI digestion and SacI + DraIII digestion, the H5 ASC & GHD was cleaved into two bands visible on the gel images (Figure 4J). The left lane of the right gel image in Figure 4J shows the undigested H5 pDZ-Rapid plasmid.

The rescued the IAV A/Indonesia/05/2005 (H5N1) virus on PR8 backbone showed clear cytopathic effect within 72 hours post-transfection however the growth kinetic curve showed a significantly reduced viral fitness relatively to the previously rescued P-Cal-P pdmH1: PR8 IAV in cell culture (Figure 4L).

### Rapid prototyping of H1 antigenic site mutants

As a proof of concept for antigen site cassette swap, the assembled PR8 H1 variant pDZ-Rapid plasmid was used to prototype an example of a PR8 H1N1 antigenic site mutant, i.e, the S12 virus (Das et al., 2013). Using the PR8 ASC mutagenesis CAD template and the published S12 HA sequence (Das et al., 2013), we designed and synthesized the ASC of the S12 virus and cloned it into the PR8 pDZ-Rapid plasmid after digestion with HpaI & BstBI (Figure 5A). Similarly using HpaI & BstBI restrictions sites, we used the P-Cal-P pdmH1 ASC CAD template to design and clone a “Delta5” mutant, where all five canonical antigenic sites were substituted with their H5 counterpart (A/Anhui/01/2005 (H5N1)) mutated to escape binding of native anti-pdmH1 antibodies (S. T. H. Liu et al., 2018; Martínez et al., 2024) (Figure 5B).

**Figure 5.**
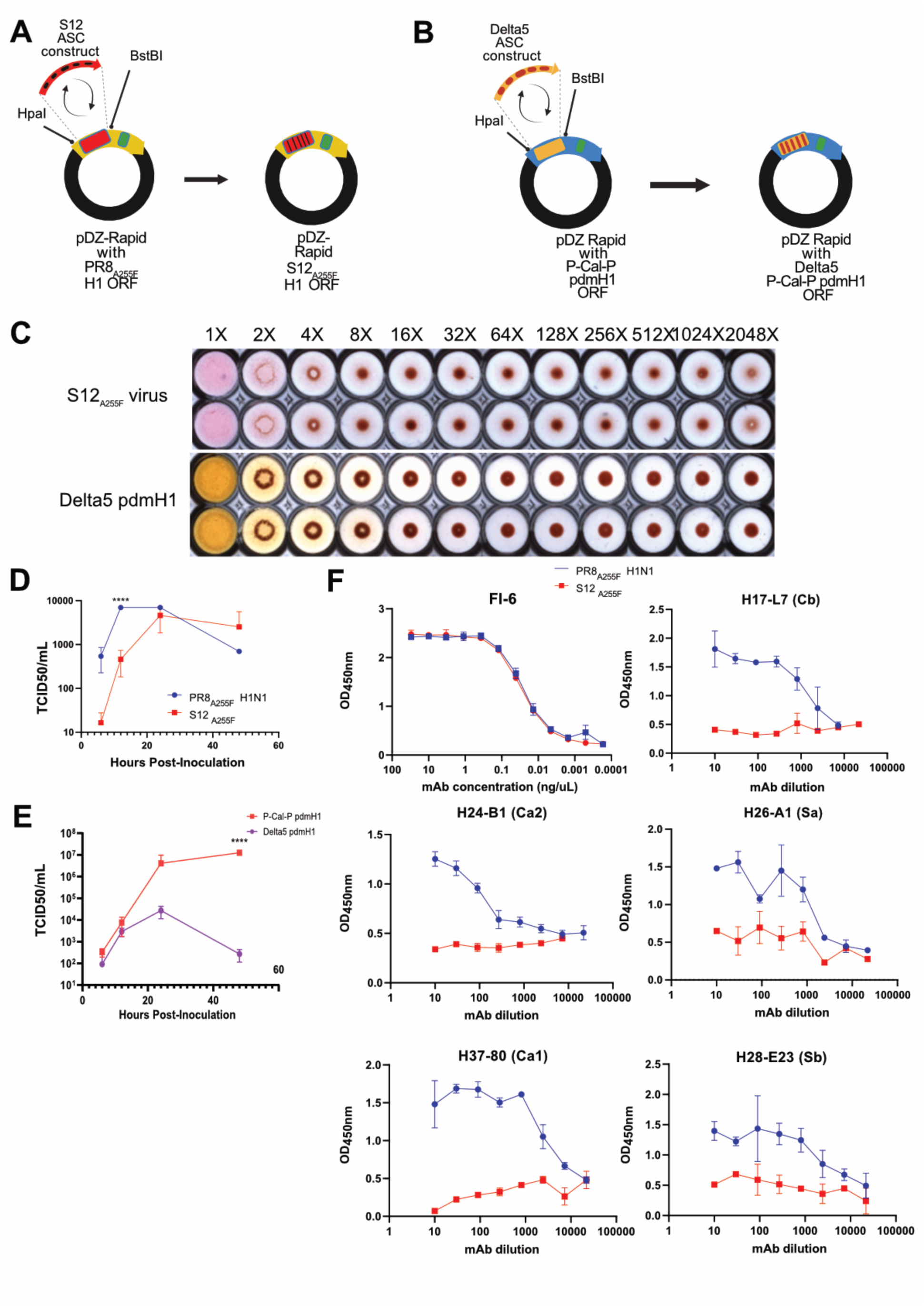
Prototyping of antigenic site mutants of PR8 and P-Cal-P pdmH1 IAVs via Antigenic Site Cassette (ASC) mutagenesis. A) Cloning of the S12 ASC into pDZ-Rapid PR8 plasmid and of the B) Delta5 P-Cal-P pdmH1 ASC into pDZ-Rapid P-Cal-P pdmH1 plasmid respectively, both linearized with HpaI + BstBI digestion. C) Hemagglutination assay of the rescued viruses. D) Growth curve characterization of rescued PR8 IAV (blue) and its S12 mutant (red) with MOI 0.01. Two-way ANOVA multiple comparison **** p-value < 0.0001 at 12 hours post-inoculation. E) Growth curve characterization of the rescued Delta5 P-Cal-P pdmH1 virus rescued on PR8 backbone, compared with growth curve of P-Cal-P pdmH1 shown earlier in Figure 3B. Two-way ANOVA multiple comparison **** p-value < 0.0001 at 48 hours post-inoculation. F) Antigenic characterization of the rescued PR8 & S12 viruses with hFI-6 mAb and a panel of mAbs specific to each of the canonical antigenic sites of PR8 H1.

Via reverse genetics, we could rescue both viruses as shown by multi-step growth curves and their ability to mediate hemagglutination (Figure 5C-E). As expected, the Delta5 pdmH1 and the S12 mutants showed significantly reduced fitness to their wild type counterparts, as their canonical antigenic sites have been drastically mutated (Figure 5D-E).

We went on to antigenically characterize the rescued PR8 and its S12 mutant a panel of five anti-PR8 HA mAbs, each specific for one of the five canonical HA antigenic sites (Gerhard et al., 1981) (Figure 5F). As expected, each mAb bound well to PR8 but failed to recognize the S12 variant. The stalk binding mAb hFI-6 (Corti et al., 2011) bound to both viruses, thus confirming proper HA folding (Figure 5F). Overall, both rescued the PR8 virus and it’s S12 mutant displayed the predicted antigenicity.

### Rapid Prototyping of H5/pdmH1 Head/stalk chimera

Finally, to prototype the H5/pdmH1 head/stalk chimera IAV, we grafted the GHD of H5 onto the GHD of pdmH1N1 using the pdmH1N1 GHD mutagenesis CAD template. In the GHD mutagenesis CAD template, we defined the HA GHD as the region bound by a conserved disulfide bond between Cys52 and Cys277 (H3 numbering) as previously described by Hai et al *(2012)*. This ubiquitious HA GHD is known to be capable of independent folding and can be easily exchanged from one HA subtype to another *(Ferrara & Temperton, 2017; Hai et al., 2012; Pica et al., 2012; Steel et al., 2010)*

To this end, we used the HpaI and MscI sites to remove the GHD of P-Cal-P H1 and replaced it with the synthesized H5 GHD of IAV A/Anhui/01/2005 (H5N1) *(Shu et al., 2006)* (Figure 6A-B). We successfully rescued the H5/pdmH1 head/stalk chimera IAV without the need for blind passaging, and observed cytopathic effect within 72 hours post-transfection. The rescued virus was able to hemagglutinate (Figure 6C), and we observed reduced virus fitness comparable to the earlier rescued IAV A/Indonesia/05/2005 (H5N1) (Figure 6D).

**Figure 6.**
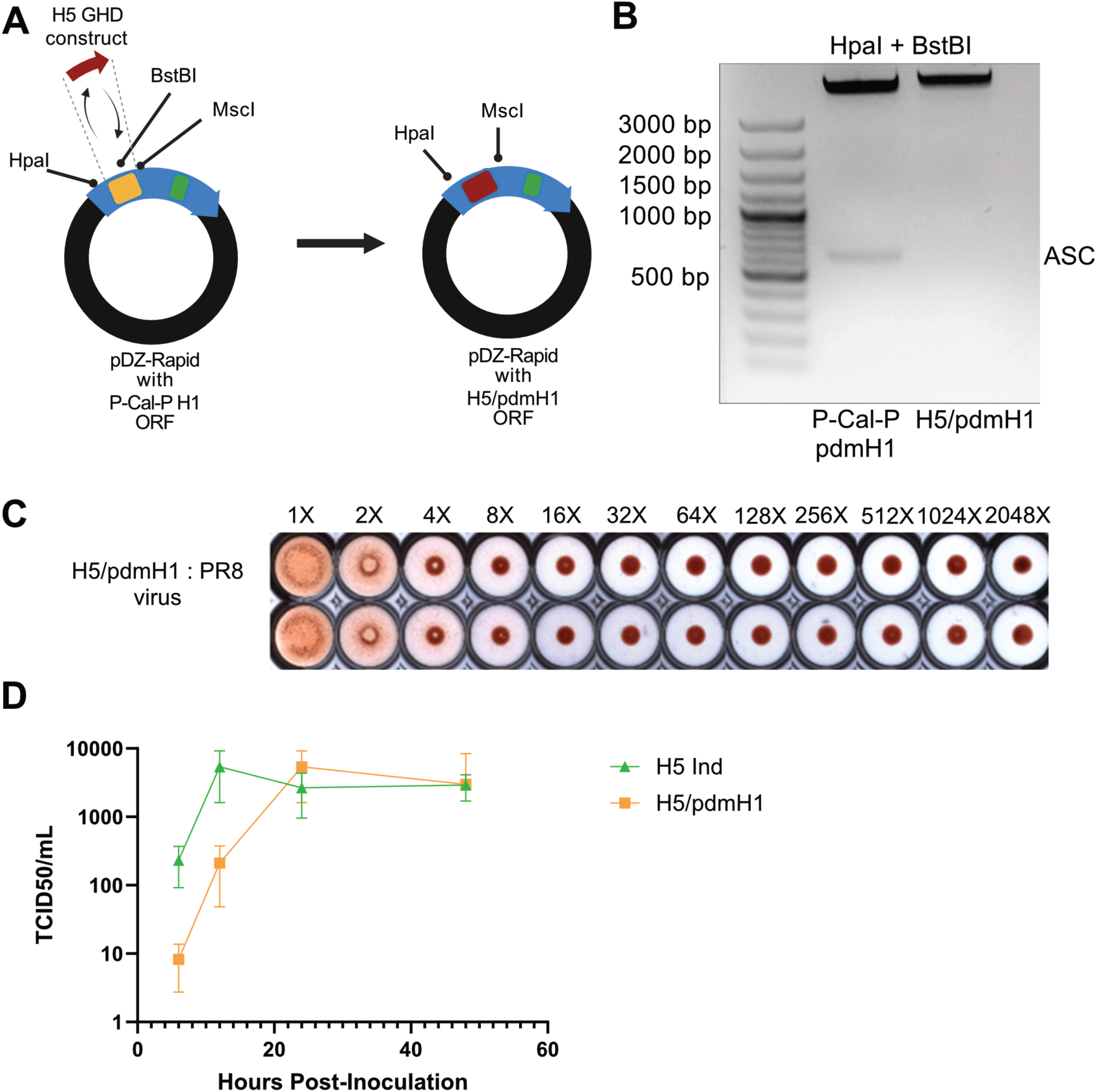
Prototyping of H5/pdmH1 head stalk chimera of P-Cal-P pdmH1 via Globular Head Domain (GHD) mutagenesis. A) P-Cal-P pdmH1’s native GHD was replaced with a synthesized GHD construct of A/Anhui/01/2005 (H5N1) via double digest with HpaI + MscI restriction enzymes. B) Gel image of restriction digest with HpaI + BstBI of pDZ Rapid P-Cal-P and pDZ Rapid H5/pdmH1. C) Hemagglutination assay of the H5/pdmH1 IAV rescued on PR8 backbone. D) Growth curve characterization of the rescued H5/pdmH1 head/stalk chimera IAV, in comparison with the previously shown growth curve of H5 Ind in Figure 4L. Two-way ANOVA multiple comparison analysis showed no statistically significant difference.

### Antigenicity testing of the H5 Virus Panel

In our study, we assembled a H5 virus panel that may be useful for studying anti-IAV humoral immune response, consisting of the attenuated IAV A/Indonesia/05/2005 (H5N1) (H5 Ind), the H5/pdmH1 head stalk chimeric IAV and the Delta5 mutant of P-Cal-P pdmH1 IAV. Our H5 virus panel was characterized for binding to hFI-6 mAb, AT225.16.69, i.e., an anti-H5 mAb, specific to IAV A/Anhui/01/2005 (H5N1) *(International Reagent Resource, 2018)*, and pooled ferret polyclonal serum raised against IAV A/Anhui/01/2005(H5N1)-PR8-IBCDC-RG6 *(International Reagent Resource, 2018a)* (Figure 7). FI6 bound to all viruses equally (Figure 7A), confirming our manipulations i.e., the antigenic site grafting, GHD grafting and deletion of the polybasic cleavage site did not cause misfolding of the synthetic HA on virions. The ferret anti-H5 (Anhui) serum and AT225.16.19 mAb showed a similar pattern with stronger binding to Anhui-derived viruses, demonstrating that the grafted H5 (Anhui) GHD folded correctly in the rescued head/stalk chimera IAV (Figure 7B-C).

**Figure 7.**
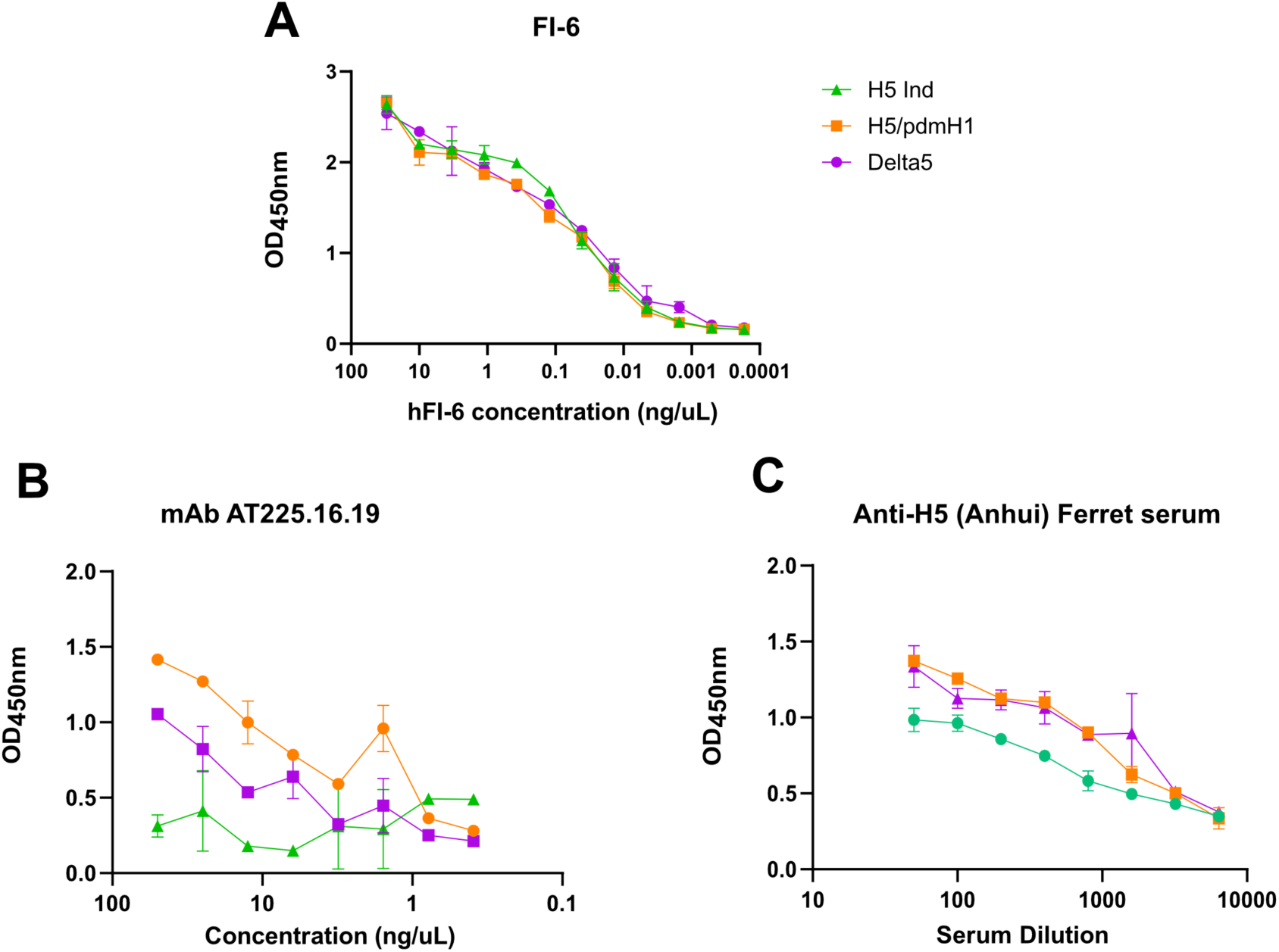
Antigenic characterization of the H5 panel. A) Normalization of H5 panel with hFI-6 mAb. The normalized H5 panel was tested with the B) AT225.16.19 anti-H5 mAb and C) ferret antiserum, both raised against A/Anhui/01/2005 (H5N1).

In conclusion, we demonstrated that by using the pDZ-Rapid plasmid we could quickly prototype and rescue an array of synthetic IAV panels for immunogenicity studies, without requiring any physical template, but only with computationally designed viral HA ORF sequences as starting material.

## Discussion

Our plasmid-based synthetic virology platform gave us the ability to manipulate IAV HA using CAD. We based our platform on modifications of the common ambisense pDZ plasmid, as it’s already a well-established and reliable system for rescue of both human and hybrid avian IAV. The strengths of our platform lie in i) Rapid CAD-assisted prototyping of synthetic IAVs with complete freedom of design ii) Modularity via exploitation of HA subtype-specific restriction site motifs that are conserved and strategically placed/inserted.

The CAD approach conferred freedom of design to prototype synthetic IAVs without needing a physical DNA template or involving complex assembly steps. For example, Ursin et al. (2022) employed PCR based site-directed mutagenesis to prototype IAV mutants with multi-antigenic site HA mutations to dissect humoral immunodominance. However, these multi-antigenic site IAV mutants are rather simplistic in design and carry no more than three-point mutations in 1-3 antigenic sites. To generate IAV head-stalk chimera, Hai et al. (2012) PCR amplified segments of different HA subtypes using primers with SapI restriction sites and cloned them into SapI digested pDZ plasmid via multi-segmental ligation reactions to assemble the final chimeric HA construct. Expanding this method, S. T. H. Liu et al. (2018) generated Δ1 mutants of Influenza A/Michigan/45/2015 (pdmH1N1) by grafting antigenic sites from H5 and H13 subtypes. Assembling the final chimeric HA constructs using multi-segment ligation reactions can be technically challenging, especially with 3 or more inserts. Furthermore, this approach necessitates each time the deconstruction and re-assembly of the entire HA ORF using multiple PCR amplified or short synthetic HA ORF segments containing the desired mutations (Hai et al., 2012; Kong et al., 2021; S. T. H. Liu et al., 2018; Martínez et al., 2024). Alternatively, CRISPR-Cas9 enables mutagenesis without deconstructing the entire HA ORF; however, this technique necessitates the availability of a physical DNA template and purified Cas9 proteins (Chen et al., 2022). Other conventional mutagenesis techniques such as Kunkel, PFunkel and QuikChange require uracil containing ssDNA templates or pre-existing DNA constructs as substrate for modifications (Firnberg & Ostermeier, 2012; Kunkel, 1985; Wrenbeck et al., 2016). Such physical DNA templates commonly originate directly from clinical or environmental isolates; however complete ORFs or individual DNA fragments can also be synthesized from available sequence information as substrates for mutagenesis.

Our CAD-based strategy therefore differs not simply in eliminating the requirements for virus-derived starting material, but in allowing the desired viral segment to be designed in silico directly from sequence data before physical synthesis (Zhou et al., 2009). Consequently, sequence information from a novel IAV isolate can serve as a starting point for the construction of a customized synthetic HA ORF, e.g. an in silico attenuated form of the novel IAV. This provides freedom of design to not only construct synthetic HA ORFs but also of the flanking 5’ and 3’ vRNA UTRs, including regulatory or packaging associated sequences (Desselberger et al., 1980) and enables the deliberate incorporation of synthetic genetic parts such as shRNA, miRNA target sites, oligonucleotide barcodes, etc. (Figure 2a) (Harding et al., 2019).

This strategy becomes conducive for synthetic vaccinology applications (Dormitzer et al., 2013) and serves as a bridge between computational antigen design and physical vaccine construction. IAV carrying synthetic vaccine antigens designed via algorithms such as consensus, COBRA (Carlock & Ross, 2023; Dzimianski et al., 2023), Epigraph (Theiler et al., 2016; Theiler & Korber, 2018), Mosaic (Fischer et al., 2007; X. Liu et al., 2023), ancestral, epitope-focused, structure-guided and generative AI protein design can be rapidly prototyped and characterized.

We identified the presence of sequence motifs in HA ORFs of various subtypes which are conserved at the amino acid level and carries strategically placed restriction sites that we exploited for cassette mutagenesis of the IAV HA ASC and GHD. These restriction sites were also specifically chosen/inserted as their restriction enzymes have versions optimized for shorter digestion time with reduced star activity were available, have several isoschizomer enzymes available from different manufacturers, and the specific combinations of these restriction enzymes (Table 1) are compatible for efficient double digest. We also cognize this restriction site-based cassette mutagenesis strategy could be scaled up to construct IAV codon-mutant libraries for mutational antigenic profiling studies (Burton & Eyre, 2021; Lee et al., 2018). For example, to study drifting antigenic sites of IAV HA, saturation mutant libraries can be prototyped via cassette mutagenesis using synthetic HA ASCs with degenerate codons e.g., NNN, NNK and NNS (Pines et al., 2022) at multiple antigenic sites of interest simultaneously (Hwang & Bang, 2016; Kong et al., 2021; Öling et al., 2018; Plesa et al., 2018; Shimko et al.,2020). Additionally, the unique BbvCI restriction site we inserted allows targeted small-scale saturation mutagenesis valuable for studying conserved regions of IAV HA.

**Table 1.**
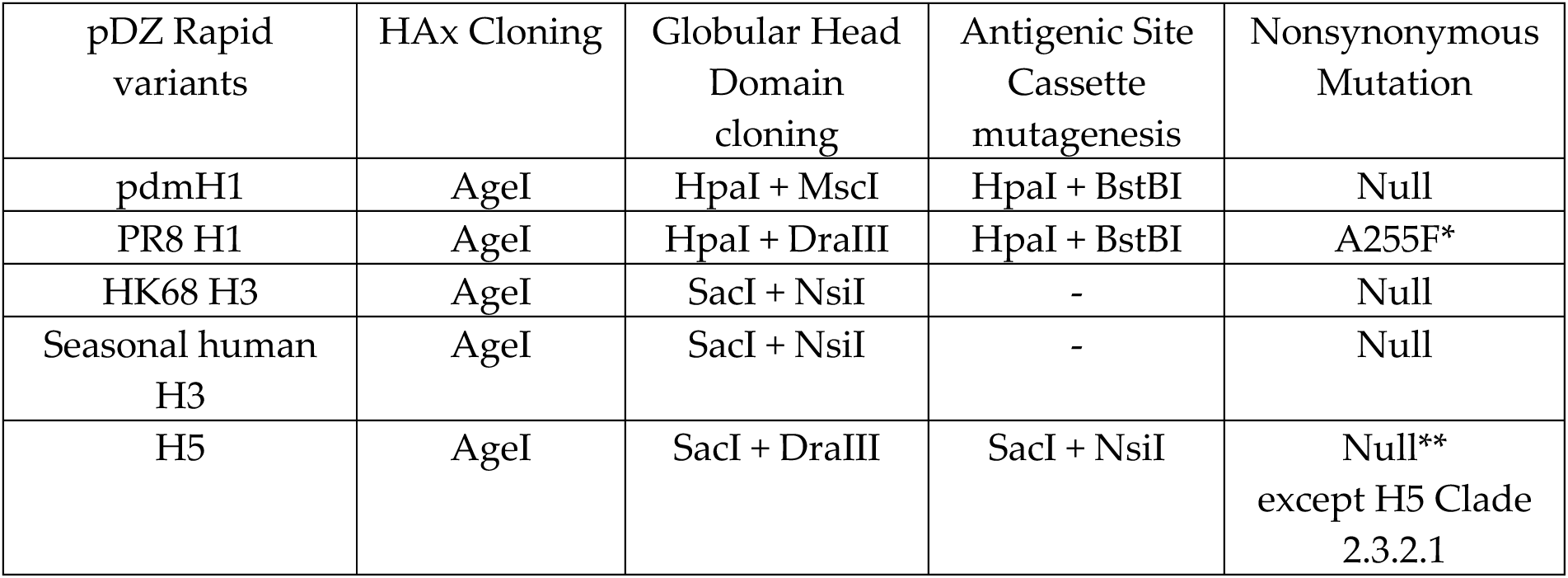
Summary of restriction enzyme double digestion combinations. *Non-synonymous mutation (based on the H1 numbering system) for the insertion of BstBI restriction site into PR8 H1 ORF. ** R277K (H5 numbering) mutation required for insertion of DraIII site for H5 Clade 2.3.2.1 (Figure S4 & S5b)

Both freedom of design and modularity confers practicality to rapidly prototype IAV candidate vaccine viruses (CVV), especially with recent public health concerns over the spillover and circulation of Highly Pathogenic Avian Influenza virus (HPAI) H5N1 clade 2.3.4.4b in the North American cattle population (Chrzastek et al., 2025; L. Wang et al., 2026; Xie et al., 2025). Even though our restriction site conservation analysis showed H5N1 clade 2.3.4.4b having well conserved DraIII restriction sites (Figure S4), we find the current CVV for clade 2.3.4.4b i.e., A/Astrakhan/3212/2020 (H5N8) (Gao et al., 2022, 2024; Pyankova et al., 2021), lacks the DraIII restriction site. Using the CAD approach to develop and characterize HPAI H5N1 CVV for rapid pandemic response, this DraIII site can be synonymously inserted during the initial design step together with multiple other manipulations of the HPAI CVV’s HA ORF, for example, performing in silico attenuation and antigenic matching of the CVV with new variants of concern through the cassette mutagenesis feature. Additionally, synthetic antigenic site mutants of the HPAI CVV can be rapidly prototyped using the cassette mutagenesis feature to investigate potential immune escape against the CVV.

From a synthetic vaccinology perspective, the modularity conferred by the cassette mutagenesis features becomes useful to rapidly prototype synthetic viral mutants & mutant libraries to study immune responses generated by the computationally designed synthetic vaccine antigens described earlier.

Our synthetic virology approach is limited by difficulty in manipulating GC-rich regions. We were unable to add any useful feature to Fragment III, which carries the species-specific RNAP I promoter (Figure 2a). The GC-rich Fragment III not only encumbered incorporation of strategic restriction sites for the removal of an RNAP I promoter containing fragment but further hindered cassette mutagenesis, as synthesizing and cloning such GC-rich RNAP I promoter cassettes are highly inefficient. Otherwise, the ability to rapidly synthesize and test different species-specific RNAP I promoters for the efficient rescue of IAVs in various cell culture systems would have been a desirable feature (Heix & Grummt, 1995; Jang et al., 2025; Kessler et al., 2024; Lu et al., 2016, 2022; Massin et al., 2005; Sato et al., 2019; Seibert et al., 2021; Suphaphiphat et al., 2010; K. Wang et al., 2017; Z. Wang & Duke, 2007).

## Conclusion

We present a versatile tool based on the improvisation of an existing ambisense plasmid, to enable CAD-assisted rapid prototyping of synthetic IAVs and to reduce scale-up technicalities in generating complex IAV mutants and mutant libraries. This tool aims to accelerate basic immunovirology research and IAV pandemic preparedness as the same plasmid can be used to simultaneously design IAV vaccine antigens and the virion tools needed to characterize their immunogenicity.

## Materials and Method

### Cell lines

HEK293T (American Type Culture Collection, ATCC, CRL-3216), MDCK (ATCC, CCL-34), MDCK cells were cultured in DMEM (Gibco, C11995500BT) with 10% (vol: vol) FBS (Gibco, 10,270–106) and 1% penicillin-streptomycin (Gibco, 15,140,163). All the cells were cultured and maintained at 37°C with 5% CO2.

### Plasmids

Influenza A backbone segment plasmids pDZ-PR8 NA, pDZ-HK68 N2, pDZ-PR8 NP, pDZ-PR8 M, pDZ-PR8 NS, pDZ-PR8 PA, pDZ-PR8 PB1, pDZ-PR8 PB2 (Martínez-Sobrido & García-Sastre, 2010) were transformed using Ultracompetent XL10 Gold E coli (Agilent). Transfection grade plasmid preparations (1 µg/µL) were made using PureLink HiPure Plasmid Midiprep kit (Thermofisher Scientific, catalog number K210004).

### Computational analyses

The GC content analysis and heatmap plot for the combined construct of Fragment I, II and III (Figure 2C) was performed with window size = 50 bp & step size = 1 bp, using a custom Python script. The code is available in Github (https://github.com/angelettilab/pDzRapid).

The conservation analysis of restriction sites on HA nucleotide sequences involved retrieval of HA nucleotide sequences, multiple sequence alignment, generation of nucleotide and translated amino acid consensus sequences, calculation and plotting of nucleotide and amino acid conservation scores.

IAV HA nucleotide sequences of H1, H3 & H5 subtypes were retrieved from the EpiFlu^TM^ database of GISAID. The searches were done by selecting the “type A”, the subtype of “H” and “N” of interest, and a particular “clade” and geographical “location” was selected only if it was of interest. Only complete original HA sequences of IAV isolates were retrieved, and HA sequences of passaged IAVs were excluded.

The EpiFlu^TM^ database has a bias, i.e., there are exponentially more sequences of IAVs isolated after 2010 especially in the case of IAV H1N1, and much fewer sequences of IAV isolated prior to the 2009 pdmH1N1 pandemic. We performed stratified random samplings to reduce this bias for IAV H1N1 sequences. We stratified the H1N1 IAV HA sequences into bins of H1N1 isolates of 1991-2010 (877 sequences), H1N1 isolates of January 2010 - December 2019 (19000 sequences), H1N1 isolates of December 2019 – August 2023 (19000 sequences), H1N1 isolates of August 2023 – May 2024 (19000 sequences), H1N1 isolates of May 2024 – January 2025 (19000 sequences), H1N1 isolates of January 2025 – November 2025 (3000 sequences). A random subset of 1000 HA sequences were sampled from each bin of H1N1 isolates, without replacement, using Python’s random.sample function with fixed random seed = 42 to ensure reproducibility. The above mentioned stratified and randomly sampled subsets of HA sequences, including all 877 sequences of H1N1 isolates of 1991 – 2010 were pooled into a single FASTA file for downstream analysis i.e., multiple sequence alignment, and restriction site conservation analysis.

Restriction site conservation analysis of NsiI site in H3N2 IAV performed on 102355 H3 sequences of human, swine, and canine lineages from 2001 to 2025.

The H5 NsiI and DraIII restriction site conservation analysis was performed using 18391 H5 sequences of North American and European H5N1 clade 2.3.4.4b IAV (Eurasian goose/Guangdong lineage). While for the H5N1 Clade 2.3.2.1, the analysis was performed using 809 sequences.

The multiple sequence alignment of the retrieved IAV HA sequences was performed using MAFFT version 7 (Katoh & Standley, 2013), using default parameters. We used a version of MAFFT 7 that is optimized for “large number of short and highly similar sequences”, which fits our purpose to analyze large number of highly similar IAV HA sequences of the same lineage or subtype.

The aligned HA nucleotide sequences were summarized into an IUPAC degenerate consensus, and restriction enzyme site was located using ambiguity-aware motif matching. A fixed-width (99 bp), codon-aligned window centered on the site was extracted from all sequences, and the consensus window was translated using an ambiguity-resolving codon expansion method. Nucleotide and amino acid conservation were computed as majority-base or majority residue frequencies, respectively. This analysis pipeline was implemented in a custom Python script which requires input of FASTA files of aligned HA nucleotide sequences, the restriction site of interest, its flanking context sequence, and the HA amino acid numbering system. The code is available in Github https://github.com/angelettilab/pDzRapid.

### Assembly of pDZ-Rapid Plasmid

A 50 µL reaction was set up to double digest 1 µg of pDZ plasmid with 1 µL of NEB EcoRI-HF (20 units) and 1 µL of NEB XhoI (20 units) in rCutSmart buffer at 37°C for 15 minutes.

Fragment I and Fragment II were synthesized as overlapping oligonucleotide fragments, without adapter sequences (Twist Bioscience). The oligonucleotide fragments had 20 bp overlap sequences with each other and with the pDZ plasmid backbone.

Fragment III was PCR amplified using Platinum SuperFii 2X MasterMix PCR reagent (Invitrogen, catalog number 12368010) according to manufacturer’s protocol and the following primers:

Forward TGT TTC TAC TAA TAA CCC GGC GGC CCA AAA TG

Reverse AAA AAG ATC TGC TAG CTC GAG CAT GCC CGG GTA CCA T

A 20 µL NEBuilder HiFi DNA assembly reaction was set up with the digested pDZ plasmid and the three oligonucleotide fragments in a 1:2:2:2 molar ratio, calculated using the InFusion molar ratio calculator. The reaction was carried out at 50°C for 15 minutes after adding 10 µL of NEBuilder HiFi DNA Assembly Mastermix. The NEBuilder HiFi DNA Assembly reaction product (2.5 µL) was used for transformation with 20 µL Ultracompetent XL10 Gold E coli. The transformants were streaked on LB Ampicillin agar plates and incubated at 37°C for 16 hours. Individual colonies were picked for further expansion in LB Ampicillin broth and mini-prepped. Sequences of the mini-prepped plasmids (Qiagen Miniprep Kit) were confirmed using NGS-based whole plasmid sequencing (Plasmidsaurus).

### Linearization of pDZ-Rapid Vector

For HAx cloning: A 50 µL reaction was set up to restriction digest 1 µg of the pDZ-Rapid vector plasmid using 1 µL NEB AgeI-HF (20 units) + 1 µL NEB recombinant Shrimp Alkaline Phosphatase (1 unit) + rCutSmart buffer at 37°C for 30 minutes.

For ASC mutagenesis of H1: The pDZ-Rapid vector was linearized using sequential digest with HpaI and BstBI restriction enzymes. A 50 µL reaction was set up to digest 1 µg of the plasmid vector first with 1 µL of NEB HpaI (5 units) in rCutSmart buffer at 37° C for 1 hour. Then 1 µL of NEB BstBI (20 units) enzyme is added to the reaction and digested further at 65°C for 15 minutes.

For Globular Head Domain mutagenesis of H1: The pDZ-Rapid vector was double digested with HpaI and MscI restriction enzymes. A 50 µL reaction was set up to digest 1 µg of the plasmid vector in rCutSmart buffer, 1 µL of NEB HpaI (5 units), 1 µL of NEB MscI (5 units) at 37°C for 1 hour.

For ASC and GHD mutagenesis of H3: The pDZ-Rapid vector was double digested with NsiI and SacI restriction enzymes. A 50 µL reaction was set up to digest 1 µg of the plasmid vector in rCutSmart buffer, 1 µL of NEB NsiI-HF (20 units), 1 µL of NEB SacI-HF (20 units) at 37°C for 30 minutes.

For ASC mutagenesis of H5: The pDZ-Rapid vector was double digested with NsiI and EcoRI restriction enzymes. A 50 µL reaction was set up to digest 1 µg of the plasmid vector in rCutSmart buffer, 1 µL of NEB NsiI-HF (20 units), 1 µL of NEB EcoRI-HF (20 units) at 37°C for 30 minutes.

For Globular Head Domain mutagenesis of H5: The pDZ-Rapid vector was double digested with DraIII and EcoRI restriction enzymes. A 50 µL reaction was set up to digest 1 µg of the plasmid vector in rCutSmart buffer, 1 µL of NEB DraIII-HF (20 units), 1 µL of NEB EcoRI-HF (20 units) at 37°C for 30 minutes.

### CAD Templates for Synthetic Oligonucleotide HA Constructs

We made annotated CAD templates for HAx ORF, epitope cassette mutagenesis, and GHD mutagenesis of P-Cal-P H1, PR8-like H1, H3 and H5 to simplify the design process. The design template sequence files can be used directly for HA sequence design and in silico simulation of the cloning reaction with the pDZ-Rapid sequence file. Sequences in the CAD templates are ready for synthesis and assembly as is provided in the supplementary.

We designed our synthetic HA oligonucleotide inserts to be flanked on both the 5’ and 3’ ends by 20 bp overlap regions complementary to the digested ends of the pDZ-Rapid plasmid. Hence, our CAD templates are compatible with both InFusion HD and NEBuilder HiFi DNA assembly seamless cloning techniques.

Snapgene software (version 7) was used to design all our synthetic HA inserts. Simulations of seamless cloning reactions using the designed HA constructs were performed to confirm functionality prior to synthesizing the oligonucleotide inserts.

### Seamless Cloning of Synthetic Oligonucleotide HA constructs into pDZ-Rapid plasmid

InFusion HD cloning was performed according to manufacturer’s protocol, and we used a vector with an insert molar ratio of 1:2 (calculated using the InFusion molar ratio calculator). A 20 µL reaction was set up using 4 µL of InFusion HD Mastermix, the linearized pDZ-Rapid plasmid (gel excised & purified with Zymoclean Gel DNA Recovery kit, catalog number D4001) and the synthesized oligonucleotide HA construct. The reaction was carried out at 55°C for 15 minutes. Upon completion, the reaction products were transformed with Ultracompetent XL10 GOLD E. coli (Agilent Technologies). The transformants were streaked on LB Ampicillin agar plates and incubated at 37°C for 16 hours. Individual colonies were picked for further expansion in LB Ampicillin broth, and transfection grade plasmid preparations were made.

### Reverse Genetics Rescue of Recombinant Influenza A Virus

A co-culture of 2 x 10^^5^ MDCK-Siat1 cells and 2 x 10^5^ HEK293T cells was prepared in each well of a 6-well plate. The co-culture was maintained overnight in DMEM + 10% (vol: vol) FBS + 1% HEPES buffer + 1% penicillin-streptomycin at 37°C with 5% CO2.

A plasmid cocktail carrying the cloned pDZ-Rapid plasmid together with pDZ plasmids carrying PR8 backbone segments (1 µg of each plasmid) was prepared in 50 µL of OptiMEM media. The TransIT-293T transfection reagent (3 µL/ µg plasmid) was diluted in 250 µL of OptiMEM media and left to incubate at room temperature for 10 minutes. The final transfection mixture was made by adding the diluted transfection reagent to the plasmid cocktail, and the mixture was left to further incubate at room temperature for 30 minutes.

The cell co-culture was gently washed with PBS before the final transfection mixture mentioned above was added. Each well was topped up with 2 mL of OptiMEM media (without antibiotics) and incubated at 37° C 5% CO**_2_** for 18 hours. The OptiMEM media was aspirated and replaced with Infection media (2.5 mL/well, DMEM + 0.3% BSA + 1% penicillin-streptomycin + 1% HEPES buffer + TPCK-Trypsin, 1 µg/µL) and incubated further at 37°C 5% CO**_2_** for 48 hours. In case of slow growing IAV mutants, the incubation was continued until visible cytopathic effect was observed.

The supernatants above (200 µL) were used to infect MDCK cell monolayers at 80% confluency. The infected MDCK cells were incubated at 37°C 5% CO**_2_** for 3-7 days. The infection passage was repeated for several rounds as described above until a marked cytopathic effect and significant viral titer was obtained.

For slowing growing IAVs, we concentrated the viruses in the supernatant via sucrose cushion ultracentrifugation prior to further passage or UV-inactivation.

Confirmation of the virus rescue was done using the hemagglutination assay as described below (Killian, 2020). Identity of the rescued virus and presence of quasi-species was confirmed via Sanger sequencing of the H**A** segment, PCR amplified and sequenced using primers:

Forward TGC AAA CTA AGA GGG GTA GCC

Reverse GCA ATC GTG GAC TGG TGT ATC T

### Sucrose Cushion Ultracentrifugation

Sucrose cushion centrifugation was performed to purify and concentrate slow-growing IAVs prior to further virus expansion, characterization, and testing. A virus supernatant sample was gently layered over 10mL of 20% sucrose in PBS in a 30mL ultracentrifugation tube. The sample was centrifuged at 131 000 g for 2 hours at 4 **°**C. The centrifuged supernatant was then decanted, and 500 µL of Hank’s Balanced Salt Solution (HBSS with Ca^2+^ & Mg^2+^, Gibco) added to the virus pellet. The pellet was incubated at 4 **°**C overnight to allow dissolution into the HBSS.

### TCID_50_ assay

Infectious titers were determined on confluent MDCK-SIAT1 cells in 96 well plates. Virus supernatants were diluted 10-fold initially and 2-fold serially diluted in infection medium and applied to cells in replicate wells. The plates were incubated for 72 hours until cytopathic effect (CPE) was evident. Wells were scored positive/negative for infection by CPE, and TCID_50_/mL values were calculated using an online TCID50 calculator of the Reed– Muench method (https://vrsxp.com/tools/tcid50-calculator/)

### Multi-step Growth curve assay

For multi-step growth kinetics, confluent MDCK-SIAT1 monolayers were infected with the rescued viruses. The supernatants were collected at 6-, 12-, 24-and 48-hour post-inoculation, clarified by brief centrifugation at 300g for 5 minutes, stored at −80 °C, and infectious titers were quantified by TCID_50_ assay. The viral titers were plotted, and two-way ANOVA multiple comparison statistical analyses were performed in Prism GraphPad version 10.1.2

### Hemagglutination assay (HA assay)

The assay was performed in a round-bottomed 96 well plate with chicken red blood cells (cRBC). The cRBC was washed with PBS and centrifugation at 800g for 10 minutes at 4 °C. The packed cRBC was then diluted in PBS to prepare a 0.5% cRBC solution A 2X serial dilution of the IAV supernatants was made up to 2^11^ dilutions with PBS. The final volume of the serially diluted IAV supernatants was 50 µL/well. The prepared 1% cRBC solution was added to the serially diluted IAV supernatants at 50 µL/well, to make a final volume of 100 µL /well. The 96 well plate was incubated at room temperature for 30 minutes prior to assessing the viral hemagglutination titer. Supernatants of slow-growing rescued IAVs were UV-inactivated and SpeedVac concentrated prior to the HA assay.

### UV-inactivation of IAV virions

One mL of the virus supernatant was added into each well of a 6-well plate, and the plate kept on ice. The UV lamp (Vilber VL-215.C, wavelength = 254 nm, power = 2 x 15 W) was placed above the 6 well plate and the virions were exposed to UV radiation for 30 minutes. The virus supernatants were collected in a glass tube and stored at -20 °C before use.

### Murine anti-serum preparation

For HK68 H3 anti-serum, C57BL7/6 mice were infected with 100 TCID_50_/mL IAV H3N2 HK68 intranasally and subsequently boosted with 100 µL UV-inactivated HK68 H3N2 intraperitoneally 1 month later.

Mice were sacrificed and serum harvested on day 3 post-boosting and inactivated at 56**°**C for 30 minutes prior to use.

### Enzyme Linked Immunosorbent Assay (ELISA)

A 96-well half area plate (Corning, product number CLS3690) was coated with 25 µL/well of 10-fold serially diluted virus supernatant at 4 °C overnight. The UV-inactivated virions were coated in duplicates. Blocking was done using 5% BSA in PBS, 50 µL/well at RT for 2 hours and the blocking solution was then flicked. The hFI-6 mAb was diluted to 4 ng/µL and 25 µL/well was added. The well plate was incubated at 37 °C for 2 hours and then washed thrice in PBS + 0.01% Tween (PBS-T). The secondary antibody i.e., HRP-conjugated goat anti-human IgG (Southern Biotech, catalog number 2040-05) was diluted 1:2000 in PBS and added 25 µL/well. The well plate was incubated at RT for 1 hour. The well plate was again washed in PBS-T thrice. TMB (3,3’,5,5’ tetramethylbenzidine, Thermo Scientific, 1-Step TMB ELISA substrate, 34029) was added 25 µL/well incubated at RT in the dark for 5 minutes before adding 25 µL/well of 2M sulphuric acid to stop the colorimetric solution. The well plate was measured for optical density at 450 nm (OD_450nm_). The results were analyzed, and non-linear regression (specific binding with Hill slope) was done in Prism GraphPad, version 10.1.2 to obtain the approximate virus supernatant dilutions required to normalize the levels of different IAV virions.

The ELISA using hFI-6 mAb was repeated by coating the well plate this time with appropriately diluted virus supernatants (dilution values obtained from the regression model) to confirm proper normalization of IAV virion levels. The post-normalization hFI-6 mAb ELISA was performed as mentioned earlier, exception the hFI-6 mAb being is 3-fold serially diluted with starting concentration of 30 ng/µL and then added 25 µL/well. If the IAV virions were not adequately normalized, we performed further normalization by calculating the area under the curve (AUC) values of the IAV virions and repeated the post-normalization ELISA until adequate normalization was achieved.

### ELISA testing of IAV virions with murine antisera/mAb

UV-inactivated virions were either normalized with hemagglutination assay or with hFI-6 mAb as above prior to coating. Half area 96-well plates were coated with 25 µL of the respective PBS-diluted virus supernatants in duplicates columns and left overnight at 4 °C. Blocking was done using 5% fat-free cow milk in PBS, 50 µL/well. The well plates were incubated at RT for 2 hours and washed in PBS-T thrice.

The following reagents were diluted in PBS-T and added 25 µL/well accordingly:

- Murine antisera were initially diluted to 1:100, and subsequently serially diluted 2-fold
- Murine anti-PR8 H1 panel mAbs were initially diluted to 1:10 and then serially diluted 3-fold
- The S-OIV 5H7 mAb (obtained from BEI Resources, NIAID, NIH: Monoclonal Anti-Influenza Virus H1 Hemagglutinin (HA), A/California/04/2009 (H1N1) pdm09, Clone S-OIV-5H7 (produced in vitro), NR-19866, Lot 70050957, 0.53 mg/mL) was 2-fold serially diluted from a starting concentration of 1060 ng/mL
- The AX-LAH3 (obtained from BEI resources, NR-50555, Lot 70029881, 1.3 mg/mL) was 2-fold serially diluted from a starting concentration of 6500 ng/mL
- The AT225.16.69 mAb (obtained through the International Reagent Resource, Influenza Division, WHO Collaborating Center for Surveillance, Epidemiology and Control of Influenza, Centers for Disease Control and Prevention, Atlanta, GA, USA. FR-527, Lot 59475327, 0.5 mg/mL) was 2-fold serially diluted with a starting concentration of 50 ng/µL.

The secondary antibody i.e., HRP-conjugated rat anti-mouse kappa light chain mAb (SouthernBiotech, 1170-05, Lot B2924-NI74E) was diluted 1:2000, added 25 µL/well and incubated at RT for 1 hour. The well plate was again washed in PBS-T thrice. 1-Step^TM^ TMB ELISA substrate solution (Thermo Fisher Scientific) was added 25 µL/well incubated at RT in the dark for 5 minutes before adding 25 µL/well of 2M sulphuric acid to stop the colorimetric reaction. The well plate was measured for optical density at 450 nm (OD_450nm_).

### ELISA testing of IAV virions with Ferret anti-sera

Half area 96-well plates were coated with 25 µL of the respective PBS-diluted virus supernatants in duplicates columns and left overnight at 4 °C. Blocking was done using 5% fat-free cow milk in PBS, 50 µL/well. The well plates were incubated at RT for 2 hours and washed in PBS-T thrice.

Pooled ferret antisera against influenza A/Anhui/01/2005(H5N1)-PR8-IBCDC-RG6 (International Reagent Resource, Influenza Division, WHO Collaborating Center for Surveillance, Epidemiology and Control of Influenza, Centers for Disease Control and Prevention, Atlanta, GA, USA, catalog no. FR-782, Lot 60267083) was serially diluted 2X from a starting dilution of 1:50.

The secondary antibody i.e., polyclonal goat anti-ferret IgG H&L (Abcam, AB112768, Lot 1082564-3, 1mg/mL) was diluted 1:500 in PBS-T, added 25 µL/well, incubated at RT for 1 hour and washed in PBS-T thrice.

The tertiary antibody i.e., HRP-conjugated polyclonal rabbit anti-goat immunoglobulin (Agilent, p0449, Lot 00006531, 0.5 g/L) diluted 1:2000 in PBS-T, added 25 µL/well, and incubated at RT for 1 hour and washed in PBS-T thrice.

The TMB color reaction was developed and stopped as described above, and the OD_450nm_ was measured accordingly.

## Abbreviations

ASC: Antigenic Site Cassette
AUC: Area under Curve
BSA: Bovine Serum Albumin
CAD: Computer-Aided Design
Cas9: CRISPR Associated Protein 9
cDNA: complementary DNA
CRISPR: Clustered Regularly Interspersed Palindromic Region
CVV: Candidate Vaccine Virus
Cys: Cysteine
DMEM: Dulbecco’s Modified Eagle’s Medium
ELISA: Enzyme-Linked Immunosorbent Assay
GC content: Guanosine-Cytosine content
GHD: Globular Head domain
HA: Hemagglutinin
HA Assay: Hemagglutination Assay
HBSS: Hank’s balanced Salt Solution
HEK293: Human Embryonic Kidney 293
HRP: Horse Radish Peroxidase
IAV: Influenza A Virus
mAb: Monoclonal Antibody
MDCK: Madin-Darby Canine Kidney
miRNA: microRNA
N: Neuraminidase
OD_450nm_: Optical Density at 450 nanometer
ORF: Open Reading Frame
PBS: Phosphate-buffered Saline
Pdm: pandemic
rHA: Recombinant Hemagglutinin
RNAP I: RNA Polymerase I
shRNA: short-hairpin RNA
S-OIV: Swine-Origin Influenza Virus
TMB: 3,3’,5, 5’-Tetramethylbenzidine
TPCK-treated: Tosyl Phenyalanyl Chloromethyl Ketone-treated
UTR: Untranslated Region
UV: Ultraviolet
vRNA: viral RNA
WT: Wild Type

## Acknowledgement

We would like to thank the International Reagent Resource, Influenza Division, WHO Collaborating Center for Surveillance, Epidemiology and Control of Influenza, Centers for Disease Control and Prevention, Atlanta, GA, USA for providing us with the anti-H5 monoclonal antibody and pooled ferret antiserum, and BEI Resources, NIAID, NIH for providing us the S-OIV-5H7 antibody to complete our study. We would also like to thank Toshimma Parris and Linnea Sjölin for having helped with the proofreading of this manuscript.

Funding was provided by Vetenskapsrådet (grant 2021-01165) and the University of Gothenburg

## Supplementary Figures

**Figure S1.**
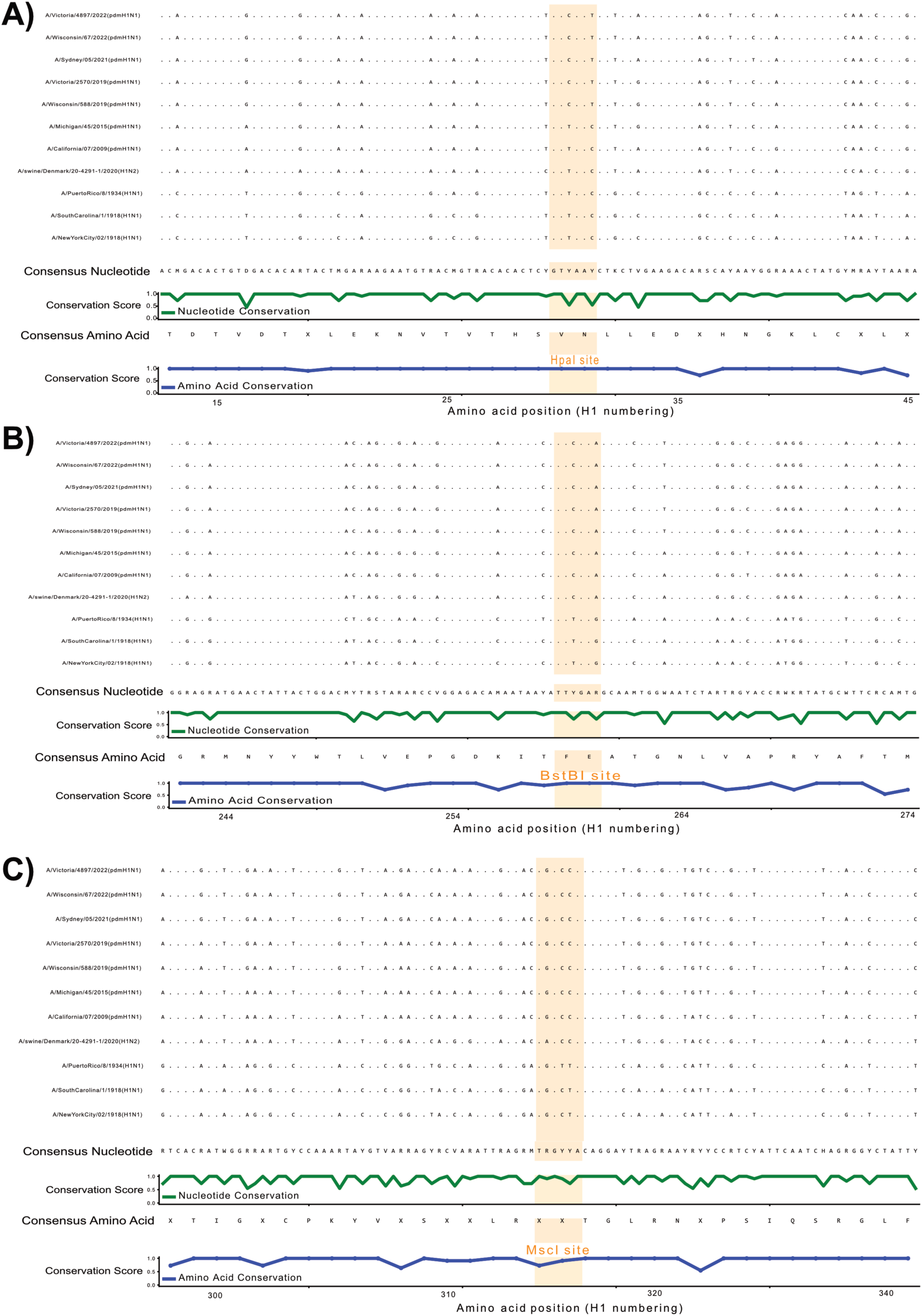
Restriction site conservation analysis of a curated set of H1 sequences. The multiple sequence alignment was performed with HA nucleotide sequences from two different H1 lineages, namely the pdmH1 and 1918 pandemic H1 lineage. The former is represented by sequences of pdmH1 seasonal IAV vaccine strains from 2009 to 2022, while the latter is represented by two 1918 H1 strains isolated in New York and South Carolina and the A/Puerto Rico/08/1934 (H1N1) strain. The H1N2 virus shown above is a special case of a pdmH1 virus which underwent reverse zoonosis into the swine population and recombined with N2 (Bhatta et al., 2020; Ryt-Hansen et al., 2025). The plot focuses on the three unique restriction sites on HA ORF of A/California/07/2009/pdmH1N1 i.e., A) HpaI, B) BstBI and C) MscI. The nucleotide (green) and amino acid (blue, H1 numbering) conservation plots show conservation at the amino acid sequences despite mutations in the viral nucleotide sequences. Hence, enabling missing restriction sites such as the HpaI restriction site (GTTAAC) to be re-inserted synonymously (GT**C**AA**T** -> GTT**A**A**C**) into post-2019 pdmH1N1 IAV strains.

**Figure S2.**
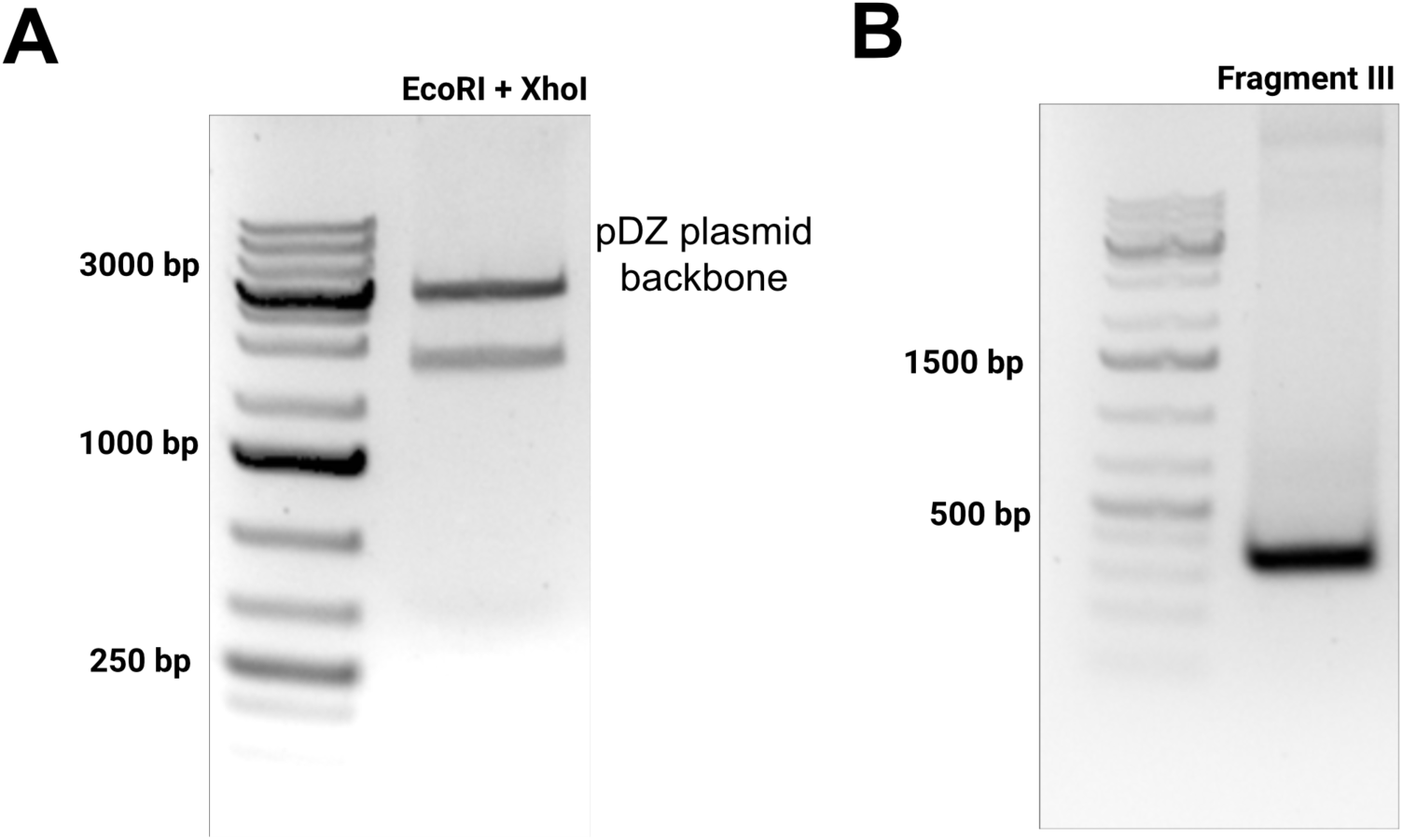
A) Digestion of pDZ plasmid with EcoRI & XhoI. The pDZ plasmid backbone seen as the upper band was gel purified and used as a vector for cloning Fragment I-III B) Fragment III PCR Amplification. Fragment III was successfully PCR amplified despite its high GC content and subsequently cloned into the pDZ plasmid backbone.

**Figure S3.**
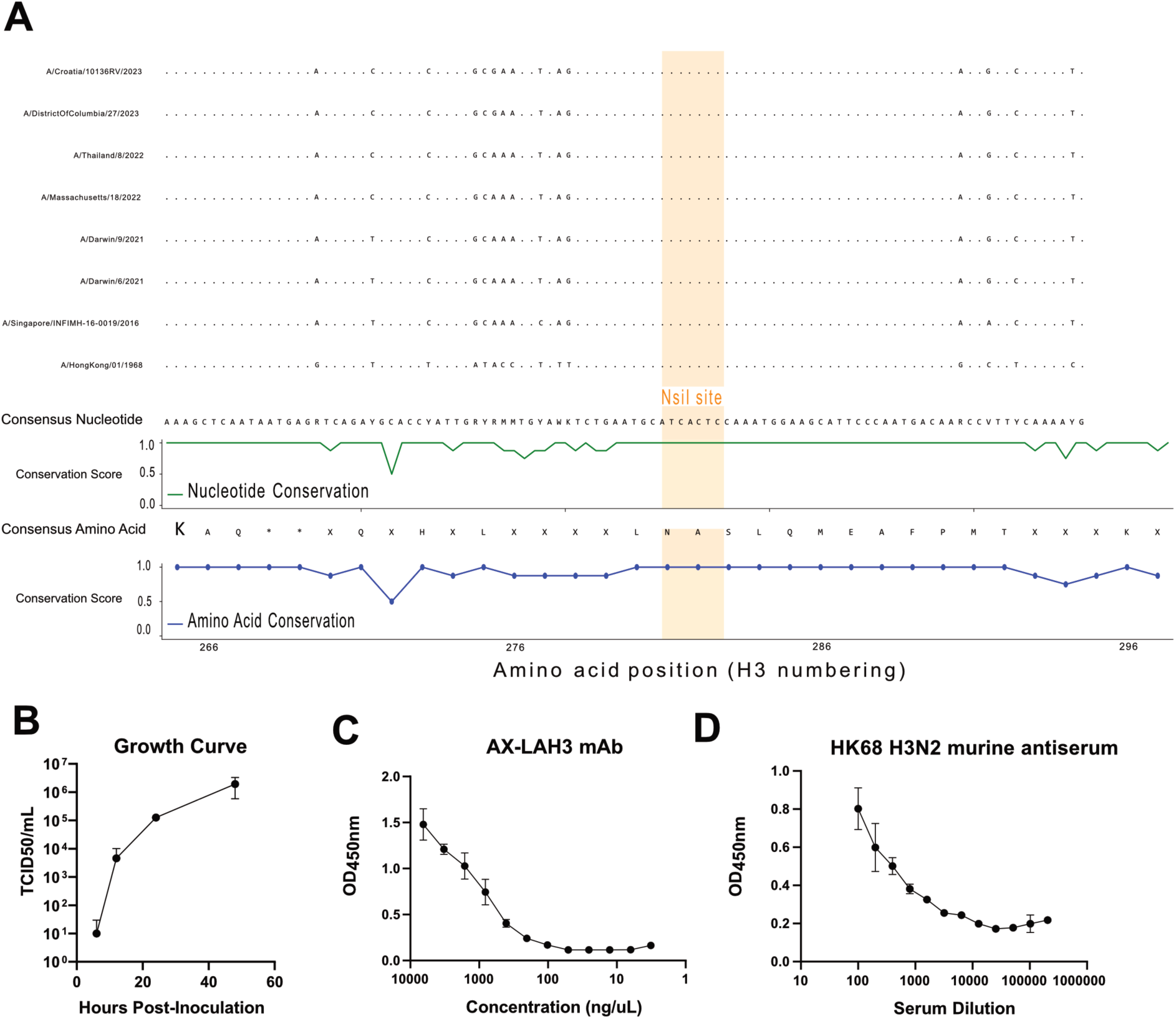
A) H3 multiple sequence alignment and NsiI restriction site conservation analysis with codon aligned nucleotide (green) and amino acid (AA, blue, H3 numbering) consensus sequences and conservation plots. The sequence alignment was performed using the HK68 H3N2 strain together with human IAV H3N2 seasonal vaccine strains from 2016 to present. Here we demonstrate that this unique NsiI restriction site (ATGCAT) in the H3 ORF has been well conserved even at the nucleotide level over the decades. Characterization of rescued HK68: PR8 H3N2 IAV. B) Growth curve of rescued HK68: PR8 H3N2 IAV with MOI 0.1. Antigenic characterization of the HK68: PR8 H3N2 UV inactivated virions by ELISA with C) AX-LAH3 pan-H3 monoclonal antibody and D) against mice antiserum immunized against wild type HK68 H3N2 IAV.

**Figure S4.**
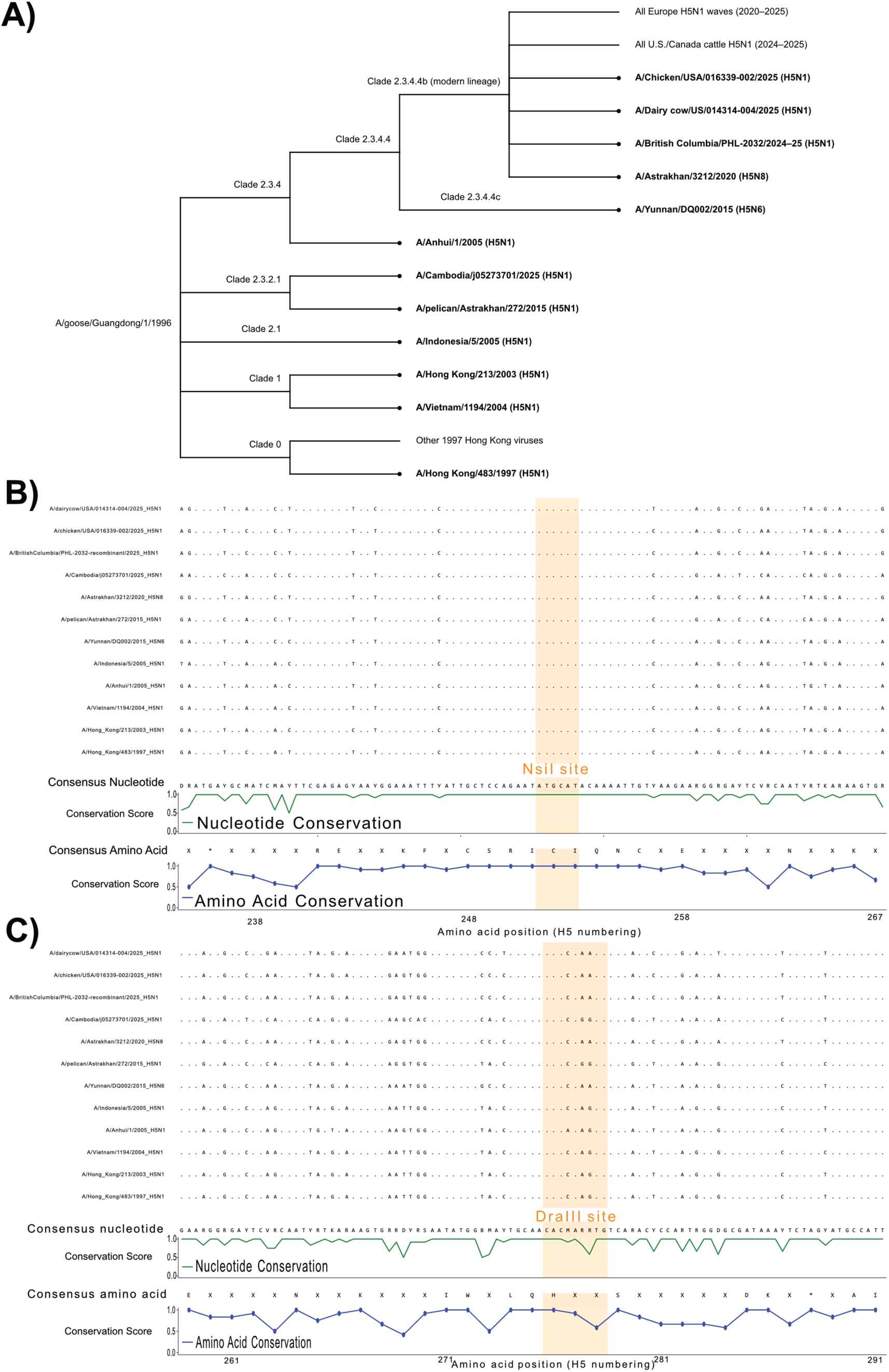
H5 phylogeny and restriction site conservation analysis with codon aligned nucleotide (green) and amino acid (blue, H5 numbering) consensus sequence and conservation plots. A) Phylogeny map for the set of H5 sequences (in bold) selected for the multiple sequence alignment and restriction site conservation analysis. The selected H5 sequences covers several clades of the Eurasian goose/Guandong lineage, ranging from clade 0 till clade 2.3.4.4b. Here we demonstrate that the B) NsiI (ATGCAT) and C) DraIII (CACNNNGTG) restriction sites in the H5 ORF has been well conserved in the H5 clade 2.3.4.4b that is currently circulating in North American cattle. We did find that certain H5 strains of clade 2.3.4.4b e.g. A/Astrakhan/3212/2020 (H5N8) lost their DraIII site. However, this restriction site can still be inserted synonymously (CACCAA**A**TG -> CACCAA**G**TG). On the other hand, H5 IAV strains of clade 2.3.2.1e such as Influenza A/Cambodia/j05273701/2025 (H5N1) requires an R277K (H5 numbering) mutation for insertion of the DraIII site to enable GHD mutagenesis (Figure S5B).

**Figure S5.**
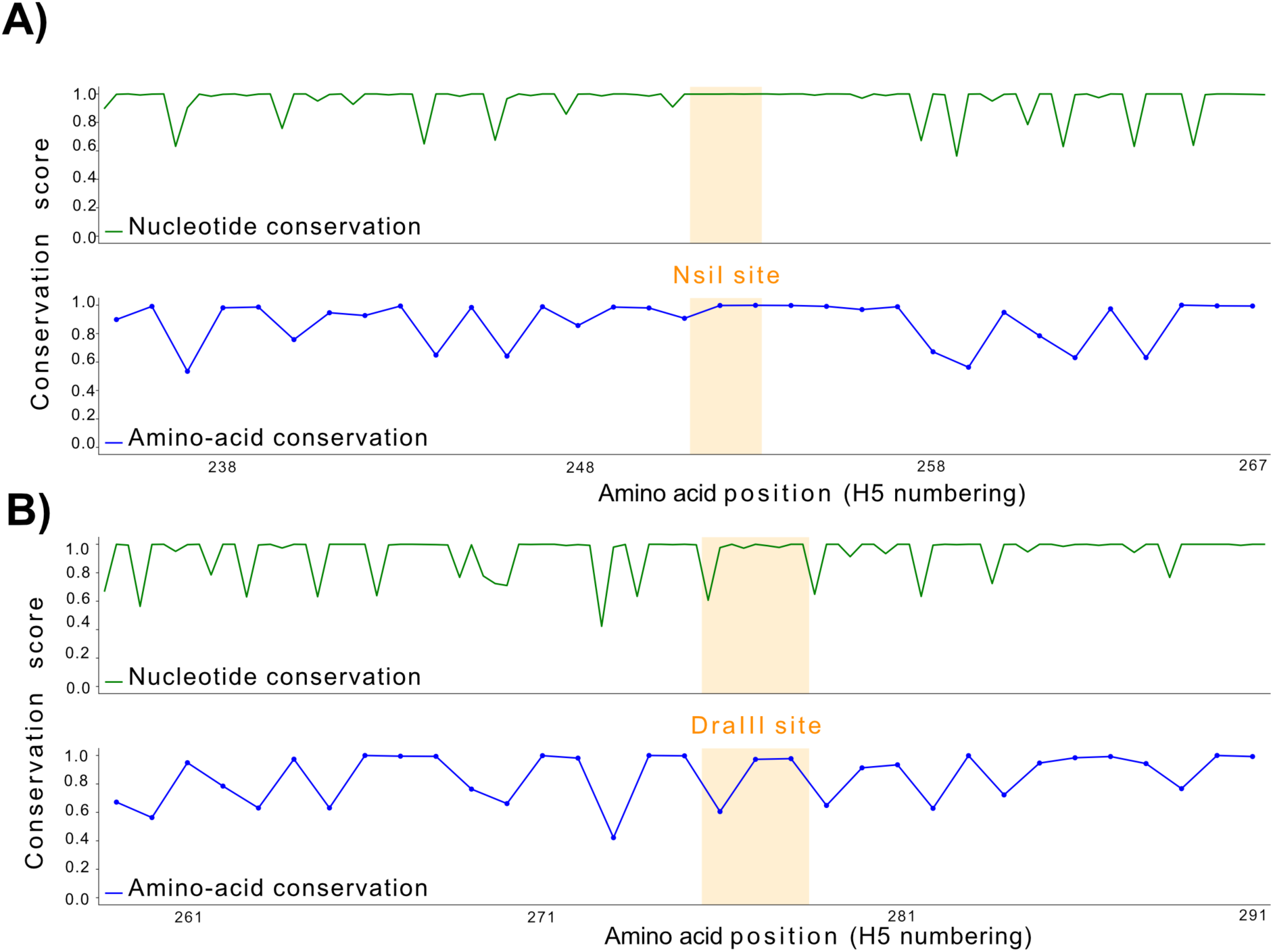
Conservation analysis of NsiI and DraIII restriction sites in H5 clade 2.3.2.1. A) The NsiI restriction site seems to be well conserved. B) The DraIII restriction site is poorly conserved with conservation scores dipping as low as 60%. Both these findings corroborate our previous findings in Figure S4 for sequences of H5 clade 2.3.2.1.

## Supplementary Files

**Supplementary File 1** CAD Template for HAx ORF cloning

**Supplementary File 2** CAD Template of Antigenic Site Cassette mutagenesis for P-Cal-P pdmH1

**Supplementary File 3** CAD Template of Globular Head Domain cloning for P-Cal-P pdmH1

